# Towards Biophysical Markers of Depression Vulnerability

**DOI:** 10.1101/2021.12.08.471836

**Authors:** D.A. Pinotsis, S. Fitzgerald, C. See, A. Sementsova, A. S. Widge

## Abstract

A major difficulty with treating psychiatric disorders is their heterogeneity: different neural causes can lead to the same phenotype. To address this, we propose describing the underlying pathophysiology in terms of interpretable, biophysical parameters of a neural model derived from the electroencephalogram. We analyzed data from a small patient cohort of patients with depression and controls. We constructed biophysical models that describe neural dynamics in a cortical network activated during a task that is used to assess depression state. We show that biophysical model parameters are biomarkers, that is, variables that allow subtyping of depression at a biological level. They yield a low dimensional, interpretable feature space that allowed description of differences between individual patients with depressive symptoms. They capture internal heterogeneity/variance of depression state and achieve significantly better classification than commonly used EEG features. Our work is a proof of concept that a combination of biophysical models and machine learning may outperform earlier approaches based on classical statistics and raw brain data.

## Introduction

Depression affects roughly one in six people^1^, and its prevalence may be increasing^2^. A major difficulty with treating depression, and psychiatric disorders in general, is their heterogeneity: a clinical phenotype or classification can arise from multiple different neural causes.^3, 4^ To address this heterogeneity, we propose describing depression in terms of interpretable, biophysical parameters of a neural model, derived from the electroencephalogram (EEG). These parameters may serve as *biomarkers*, variables that allow subtyping of depression at a biological level. They can be thought of as latent variables that may capture individual differences between patients. As a proof of concept, we show that this idea works even in a small patient cohort.

Several studies have used EEG data to identify potential biomarkers for psychiatric disorders.^5–8^ These studies emphasized EEG features: components of event-related potentials (ERPs) or oscillatory responses, not biophysical parameters. For example the paper ^6^ found smaller N1 amplitudes in depression, see also^9, 10^. However, the results from these studies are difficult to connect back to biology: The data features (e.g. ERPs, oscillatory responses, resting state activity) do not directly map back to brain structures or to physiologic changes. There are exceptions, e.g. the loudness dependence of the auditory evoked potential^11–13^, but in general these analyses are mainly phenomenological. They also fail to consider depression’s internal heterogeneity, which limits generalizability of the derived biomarkers. A recent meta-analysis suggested that no EEG marker had reliable clinical utility^14^, although some recent work has tried to address this.^15, 16^

One approach to overcoming depression’s heterogeneity might be to shift the level of analysis. For instance, source localization techniques can interpret scalp phenomena in terms of their underlying cortical generators.^8, 17^ Still these methods emphasize waves/patterns in the electrical activity whose neural basis remains unclear. A deeper level of analysis more grounded in cellular physiology may be possible when using biophysical models. This is the approach we take here using biophysical models. Their parameters describe the neurobiology or neural populations (e.g. synaptic time constants, intrinsic and extrinsic connectivity) that give rise to the scalp-recorded patterns. They capture important developmental, structural and functional properties of cortical sources. Synaptic time constants are important for determining the EEG signal.^18^ Intrinsic and extrinsic connectivity go through characteristic changes throughout the development of the brain and can exhibit differences with age or in the presence of a disease.^19^ For instance, we and others have used biophysical models to analyse data from patients with neurological diseases recorded using M/EEG and fMRI. ^20–24^

Here, we constructed biophysical models using Dynamic Causal Modelling (DCM). These describe the cortical network activated during a cognitive conflict task that activates depression-relevant brain areas.^25–27^ Our model transforms high-dimensional EEG data onto a mechanistically interpretable feature space^20^; in which, we show below that we can better measure depression’s internal heterogeneity. We present a proof of concept for the following idea: that biophysical model parameters yield a low dimensional, interpretable feature space. As a result of that better capture of internal heterogeneity/variance, model-derived features achieved significantly better classification than manifest EEG features. Our work shows that a combination of biophysical models and machine learning may outperform earlier approaches based on classical statistics and raw brain data.

## Materials and methods

### Dataset

The dataset included 15 psychiatric patients who reported current or past depressive symptoms and 34 non-diagnosed controls. Importantly, this dataset was not limited to patients diagnosed with unipolar depression, but included bipolar and unspecified depression. We considered this a better demonstration of our approach to heterogeneity. This is a secondary analysis of a cohort collected in a previous study.^28^ For details of the EEG recordings, see Supplementary Methods.

EEGs were collected as participants performed the Multi-Source Interference Task (MSIT), see Figure 1C.^29^ MSIT has been validated to produce robust cortical activations at the single-participant level, in both fMRI^30^ and EEG^31^ studies, and is used for assessing depression state. For more details regarding the task and dataset, see Supplementary Methods.

**Figure 1:**
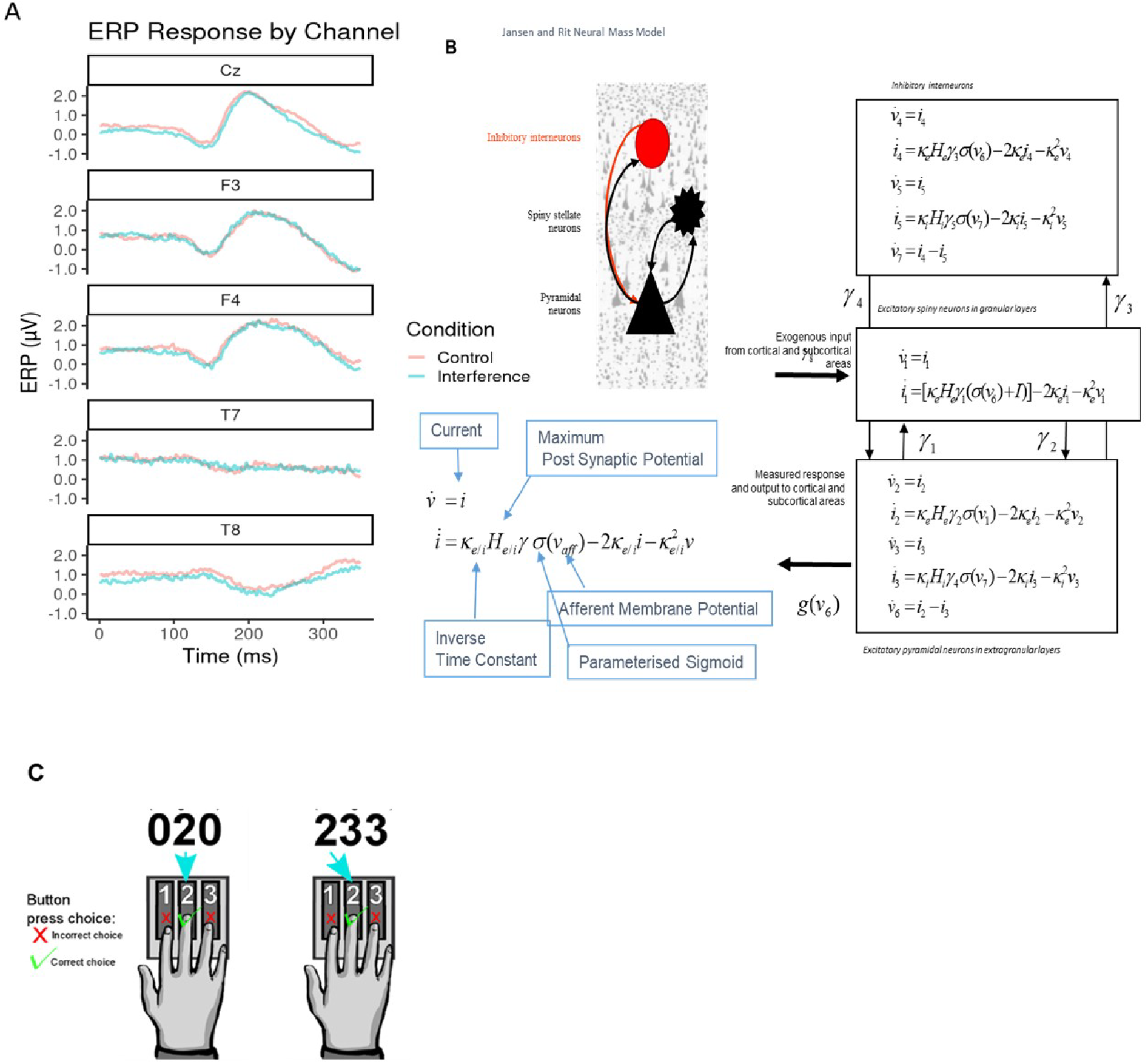
(A) Sample average EEG respouses. These are shown per channel retained for classification analysis. Both control (red) and interference condition (blue) averages are plotted across the 350ins response time post stimulus onset. Channels Cz, F3, and F4 show typical ERP amplitudes, while T7 and T8 show flat EEG responses across the diuation. **(B) Jansen and Rit Model.** The schematic diagram summarizes the evolution equations that specify a Jansen and Rit (JR) neural mass model of a single souice. This model contains three populations, each loosely associated with a specific cortical sub­population or layer1: pyramidal and spiny stellate neurons and inhibitory interueurons. Second-order differential equations mediate a linear convolution of presynaptic activity to produce postsynaptic depolarization. This depolarization gives rise to firing rates within each sub-population that provides inputs to other populations. The operations are captured by the equations shown on the right-hand side, which are explained in the bottom-left comer inset that includes the par ameters appearing in these equations and then definitions. For a thorough discussion of these equations, see the study by ^26^**.(C) Multiple Source Interference Task (MSIT).** The task requires participants to report on a presented stimulus by using their’ index, middle or ring finger to press three buttons corresponding to number’s 1, 2, and 3 respectively. The stimulus appears on a screen displaying three numbers; one number (the target) is different from the other two (distractors). The participant identifies the target by pressing the corresponding buttons. There are two task conditions, control and interference. During control trials, the distractor numbers are zeros and the location of the target number is aligned with its corresponding button. In interference trials, the distractors are non-zero numbers, and the target is hi a location misaligned with that of the button,

### ERP Analysis

P300 components were extracted from 70 EEG channels (average ERPs over participants are included in Figure 1A; see also Suppl. Figure 1). Previous MSIT studies found differences in the P300 component between task conditions.^32, 33^ P300s are a common signature of conflict and cognitive control, and arise when incongruent stimuli are processed.^34, 35^ We used P300 peak amplitude and latency as EEG classification features (see Class Balancing and Model Training section below).

### Dynamic Causal Modelling (DCM)

We used Dynamic Causal Modelling (DCM)^20–22, 36–41^ to infer processes at the neuronal level from scalp EEG measurements.^2^ We characterized changes of intrinsic (within area) and extrinsic (between area) connections across task conditions and between individuals. We assessed whether information flow changed in the same way (top-down, bottom-up or both) between the two task conditions across all participants. We here used DCM for Evoked Responses and Jansen and Rit (JR) mass model (Figure 1B). JR models can predict both evoked and induced responses and have been used in theoretical and experimental studies.^27, 43–46^ DCM was implemented using SPM12. For more details about DCM, see Supplementary Methods.

### Functional Network

The functional network modelled with DCM can be seen in Figure 2 (cf. model M1 in top left corner, all other models include the same network and assume changes in different connections, explained below). This network is comprised of areas activated during the MSIT.^29, 53^ The network included sensory, temporal, parietal, dorsal and ventral frontal areas, and ACC: V1, dACC and the following areas in both hemispheres: ITG, SPL, vlPFC, dlPFC. Changes in functional connectivity within this network were observed, at the group level, in patients with depression.^42, 54–56^ For details about the coordinates of these areas, and why we chose this network and not other areas, see Supplementary Methods. We used DCM parameter estimates as data features for classification and clustering.

**Figure 2:**
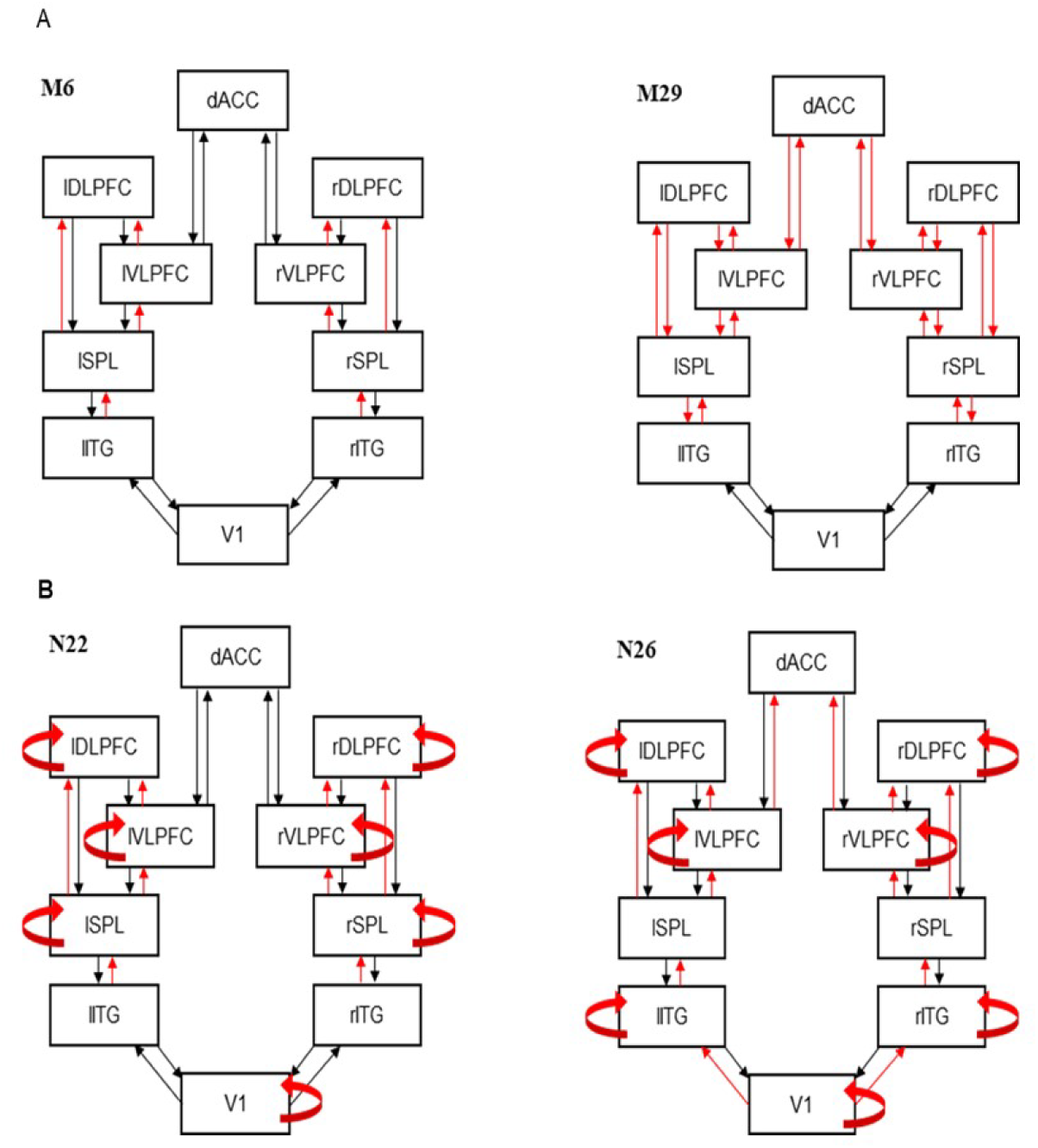
Best fitting models. (A) DCM best fitting model M6 (left) and runner up model M29 (right).Model M6 includes changes in forward connections at all levels except VI. The runner up (M29) is very similar. It also includes the corresponding feedback connections on top of the forward connections included in M6. (B) Best fitting DCM models showing modulations of intrinsic connections for controls (N22; left) and patients (N26; right).

### DCM parameters

DCM parameter estimates were obtained by fitting ERPs, i.e. P300 potentials evoked during the MSIT task. We fitted ERP recordings from different participants, patients and controls. We thus obtained DCM parameter estimates. Noise or heterogeneity in the scalp-level recordings might arise from a small number of disruptions in the underlying network. After fitting, variability in ERP recordings leads to variability in the biophysical model (DCM) parameter estimates across participants. This, in turn, could describe biotypes or endophenotypes of depression. We hypothesize: 1) If DCM can capture that variability, then DCM-derived model parameters might be more effective than raw ERPs at classifying patients from controls; 2) Clustering of DCM parameters may help identify clusters of endophenotypes. We tested these hypotheses below.

DCM parameters were obtained after fitting data from individual subjects. These included the following parameters: extrinsic connectivity, *A* (12x2=24 parameters), differences in extrinsic connectivity between MSIT conditions, *B* (24 parameters, derived from the model fitting as shown in Results), excitatory and inhibitory receptor density, *G* (2x10=20 parameters), strength of connections between the three populations of the JR model shown in Figure 1B, *H* (4 parameters; see arrows in Figure 1B), and excitatory and inhibitory synaptic time constants, *T* (20 parameters).

### Model comparison

Because we did not know how connectivity changed between task conditions, we compared several variants of the biophysical model describing the network of Figure 2. We considered a network containing all of our modelled brain regions: V1, ITG, SPL, vlPFC, dlPFC, and dACC. We assumed forward and backward connections between specific areas, as well as lateral connections between homologous areas in the right and left hemispheres. We asked which connections might change between MSIT conditions and considered all possible changes. The alternative model variants differed in the connections that could change. Following ^40^, we first considered changes of extrinsic connections (i.e. between nodes) only. Then in step 2, changes in intrinsic connections. The first twenty candidate models with extrinsic connections changes that we considered, are shown in Suppl. Figure 2. Overall, the candidate model space comprised 45 models. We describe in detail these 45 models in Supplementary Methods. The 45 models included all possible models where forward or backward connections changed between different parts of the brain network. Finding the most likely among these models yielded the extrinsic connections that were modulated during the task. For model comparison, we used an approach known as Bayesian model selection (BMS). This was performed assuming fixed-effects (FFX).^57^ BMS fits competing models to EEG data and assesses the most likely model. See Supplementary Methods for more details.

We considered variations of the network shown in Figure 2. We assumed changes in intrinsic connections from each node to itself (in addition to changes in extrinsic connections that the winning model above assumed).^40^ We thus assumed that intrinsic connections could change at any (combination of) brain areas: V1, ITG, SPL, {vlPFC, dlPFC} and dACC. We thus compared 32 candidate models in total.

### Classification features

The biophysical parameters of the best DCM model obtained via BMS were used as features for patient classification and subtyping.

Two sets of classification features were used. DCM parameters and EEG features. DCM parameters were directly compared to EEG features. The DCM parameters included intrinsic and extrinsic connections that were found to differ between MSIT conditions in both the patient and control DCM fits. This resulted in 92 DCM parameters. These were used as DCM predictors in machine learning classifiers.

To compare the predictive power of the DCM parameter estimates against the EEG features, we used an equal number of ERP features (92). The full set of potential EEG features included 240 variables (60 EEG Channels x 2 conditions x 2 variables, i.e. ERP peak amplitude and latency differences between the two MSIT conditions). We reduced the number of channels to 23 so that the total number of EEG features was the same as the number of DCM parameters. This reduces bias in the comparison between ERP and DCM feature sets. To choose these 23 channels, we performed permutation testing that assesses the change in prediction error of classification after permuting a feature.^60, 61^ The ERP features were chosen based on their contribution to a random forest model (constructed without hyperparameter tuning). This “naive” random forest allowed us to select channels with features that were most beneficial in separating classes while still allowing for multiple interaction effects between features. The tradeoff of this method stems from using a reduced number of features with the benefit that they are potentially more meaningful, and easier to interpret.

Critically, the above selection of ERP features, biases the subsequent machine learning analysis *against* our a priori hypothesis that DCM-based features will provide superior classification and clustering – the DCM analysis considers an unselected set of model parameters, whereas the ERP analysis begins with features already known to have some classification power. The selected channels are included in Supplementary Table 2.

### Class Balancing and Algorithm Training

The dataset was imbalanced between control and patient classes. Only 15 of the 49 participants being patients with depressive symptoms. We implemented Synthetic Minority Over-sampling Technique (SMOTE) to correct for this imbalance.^62^ For more details, about oversampling, see Supplementary Methods. This brought parity to the classes with 34 observations each (patients and controls).

Given the sample size limitations, we used 10-fold cross-validation to train and tune machine learning classifiers.^59, 66^ See Suppl. Figure 9 for a visual depiction of the sampling strategy and Supplementary Methods. The cross-validation was used to train classifiers and assess whether DCM features can better measure depression’s internal heterogeneity, compared to EEG features.^66^ We compared the performance of different machine learning algorithms to distinguish patients with depression vulnerability from controls in both the EEG and DCM feature sets. The algorithms included Support Vector Machines (SVM)^67^, Random Forests^60, 95^, and Gradient Boosted Trees^68^. Comparing DCM and EEG features using multiple algorithms ensures that conclusions are not sensitive to the specific algorithm used. We also used multiple performance metrics, including *F*-measure (F1-score), and the Matthews’ Correlation Coefficient (MCC).^69^ See Supplementary Methods for more details.

### Feature Importance

The best performing classifier as determined by mean MCC score was used to compute feature importance. Shapley additive explanation (SHAP) values were constructed for predictions on the original data set (49 participants, no SMOTE augmentation).^72^ This reveals how efficient the low dimensional space spanned by DCM and EEG classification features is in describing the internal heterogeneity of patients with depressive symptoms. SHAP values were constructed using subsampling of different combinations of input features and attributing a weight representing how much credit features should receive for class prediction. The predictive power of EEG and DCM features was compared directly because the corresponding SHAP values take on the same scale and are predicting the same underlying data.

### Unsupervised Clustering

The ten most important features as determined by SHAP values from both the ERP and DCM feature sets were used to construct embedding scores with t-stochastic neighbor embeddings (*t-*SNEs). *t-*SNEs are useful for exploring higher-dimensional data in lower dimensional representations when non-linear relationships exist in the data.^73^ For more details on t-SNE see Supplementary Methods. This provided visualizations of the data that were convenient for assessing subtypes or clusters of patients with depressive symptoms.

Clustering was performed using *k-*means in the three dimensional space obtained by *t-*SNE. *K*-means is a unsupervised machine learning method that groups observations to reduce within-cluster sum squares distances and increase the sum squared distance between cluster centroids.^74, 75^ *K*-means depends on an *a priori* number of clusters. The optimal cluster number can be found by computing Silhouette scores across candidate values of *k*. Observations which been classified appropriately have a lower mean distance between points within their assigned cluster compared to the mean distance to points in the next-nearest cluster neighbors.^76^ This ratio is given by Silhouette scores.

## Results

### DCM

We first asked how information flow changes between the congruent and incongruent condition of the MSIT. We used Bayesian Model Selection (BMS; see Supplementary Methods) to find the connectivity pattern between our six modelled areas. We fitted ERP data using a biophysical DCM model (Figure 1B) and scored all possible model variants that corresponded to different subsets of connections that might change between MSIT conditions. Different competing models represented different combinations of modulated forward or backward extrinsic connections (see also Methods). BMS identified the winning model as M6 (*BF>3*; Figure 2A, see also Suppl. Figure 4). Model M6 includes changes in forward connections between ITG◊SPL, SPL ◊vlPFC, SPL ◊dlPFC and vlPFC◊ dLFPC. The runner up (M29) is very similar to M6 (Figure 2B). It includes the corresponding feedback connections on top of the forward connections included in M6. M29 also includes changes in feedforward and feedback connections between vlPFC, dLFPC, and dACC. Thus, the prominent difference between task conditions was changes in the forward (bottom-up) information flow between sensory processing and associative regions. This corresponds to different processing of sensory input between the two MSIT conditions that gives rise to ERP differences in the first 350ms after stimulus presentation.

In the second step, BMS was used to test the modulation of the intrinsic connections. In this step, we fixed the extrinsic connections to those shown in the winning model M6 above. We then considered variants of M6, where intrinsic connections (within each brain area) were allowed to change. These variants formed a different model space to the one considered above (Methods and Suppl. Figure 3). BMS identified that variant N22 had the highest evidence (*BF>3;* Figure 2B, see also Suppl. Figure 5). This included modulated intrinsic connections in V1, SPL, vlPFC and dlPFC areas. The runner up (N23) is exactly the same as N22 with the additional modulation of dACC intrinsic connections (Suppl. Figure 5). We also performed the same BMS for the patient cohort.

We repeated the earlier analysis, where we fitted the neural mass model to patient data only. We wanted to capture the intrinsic heterogeneity. We first found that model M8 best described the changes in extrinsic connections (Suppl. Figure 6). This included changes in all feedforward connections. The runner up (M41) was similar to the winning model, M8, in that it included changes in the feedback connections too. The difference from M8, was that it did not include changes in feedforward (and feedback) connections between V1 and SPL and vlPFC and dACC. The winning model for patients, M8, was similar to the winning model for the control cohort (M6). The difference between the winning models was that two more brain regions showed modulations of feedforward connections. Patients showed changes in vlPFC to dACC and V1 to ITG, in addition to changes in forward (bottom-up) information flow between the ITG to SPL, SPL to vlPFC and dlPFC, and vlPFC to dlPFC that we had found for controls. However, we did not use this difference for our clustering analyses below. Thus, we do not claim that the additional connection changes that we found for the patient cohort are biomarkers of depression or depressive vulnerability. Rather, the *similarities* (common connections) between M6 (controls) and M8 (patients) were the inputs to machine learning algorithms together with the biophysical parameters described in the last paragraph of this section below.

In the second step, similar to the analysis above, we identified N26 as the model with highest evidence (*BF>3*; Figure 2B, see also Suppl. Figure 7). This is very similar to N22 that was the winning model for controls. The only difference between N22 (controls) and N26 (patients) is that N22 includes changes in intrinsic connections in ITG instead of SPL. The runner up (N9) is also very similar and assumes modulations of intrinsic connections in dACC instead of V1. Again, we are not claiming this change to be a biomarker of depression, but as an example of how DCM can identify underlying neurological variability. The clustering/classification analysis used only the intrinsic connections that were common to N26 and N22.

Several connections were independently shown to be modulated between conditions in both cohorts (controls and patients): ITG to SPL, SPL to vlPFC and dlPFC, vlPFC to dlPFC, and the intrinsic connections in V1, vlPFC and dlPFC. After finding the set of extrinsic and intrinsic connections that changed between MSIT conditions, we fitted the winning model (N22 for controls; N26 for patients) to each participant. Example model predictions for each of the three populations are shown in Suppl. Figure 8. Blue and red lines correspond to the two MSIT conditions (control and interference). There are three pairs of lines, corresponding to the three populations of the JR model (cf. Figure 1, right panel). After fitting the model, we obtained connections (*A, B*) and other biophysical parameter estimates from each participant (*G, H, T*; see Methods). These were used as DCM predictors in the next section.

### Classification

Our goal was to assess whether DCM features can better measure depression’s internal heterogeneity, compared to EEG features. To do this, we asked whether DCM features achieved significantly better classification than EEG features.^66^ EEG features included ERP parameters, i.e. ERP peak amplitude and latency differences between the two MSIT conditions. We used ERPs from 24 channels so that the number of EEG predictors was equal to the number of DCM predictors. This allowed us to perform head-to-head comparisons between ERP and DCM predictors. Crucially, we chose ERP predictors in such a way that it biases subsequent analysis against DCM predictors (they were the best classifiers in an initial random forest model; see Methods for more details). Due to the small sample (15 patients), it was not possible to test out of sample predictions for algorithm robustness. To assess the relative ability of DCM and ERP parameters to capture diagnostic heterogeneity (and thus support better classification), we computed classification performance using three algorithms: SVM, Gradient Boosted Tree and Random Forest (Figure 3). The SVM algorithm performed best for both the DCM and ERP data sets (highest average MCC scores).

**Figure 3:**
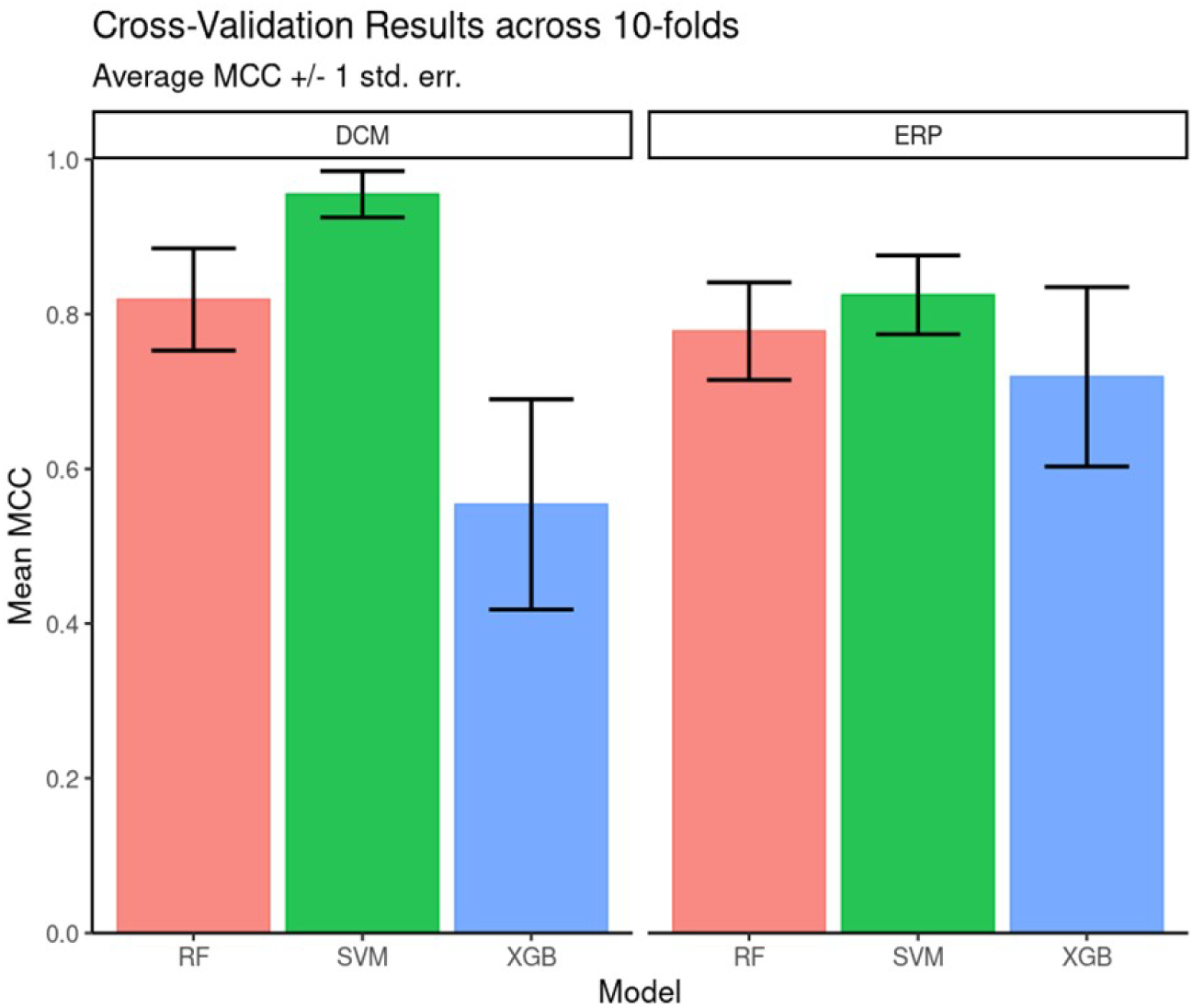
Cross validation performance. Matthew’s. Correlation Coefficient scores across the 10-fold cross-validation. Mean and +/- 1 standard error are reported for each model architecture. SVM classifiers performed the best for both feature sets on average across the 10 folds.

Overall, DCM features led to better classification accuracy than EEG features across all three algorithms tested. SVM was used for assessing feature importance because it had the highest classification performance (see Suppl. Table 3 for full results of MCC and F-Score). It also had a more parsimonious hyperparameter set (two – kernel gamma and cost) compared to either decision-tree ensemble model.

To evaluate feature importance, we used absolute SHAP values averaged across participants. Figure 4 shows SHAP values for the 10 most important DCM (Figure 4A) and EEG (Figure 4B) features. By taking the average across each of the 49 classification decisions (participants), we have a relative rank of feature importance. Overall, the DCM features have higher mean absolute SHAP values compared to EEG features.

**Figure 4:**
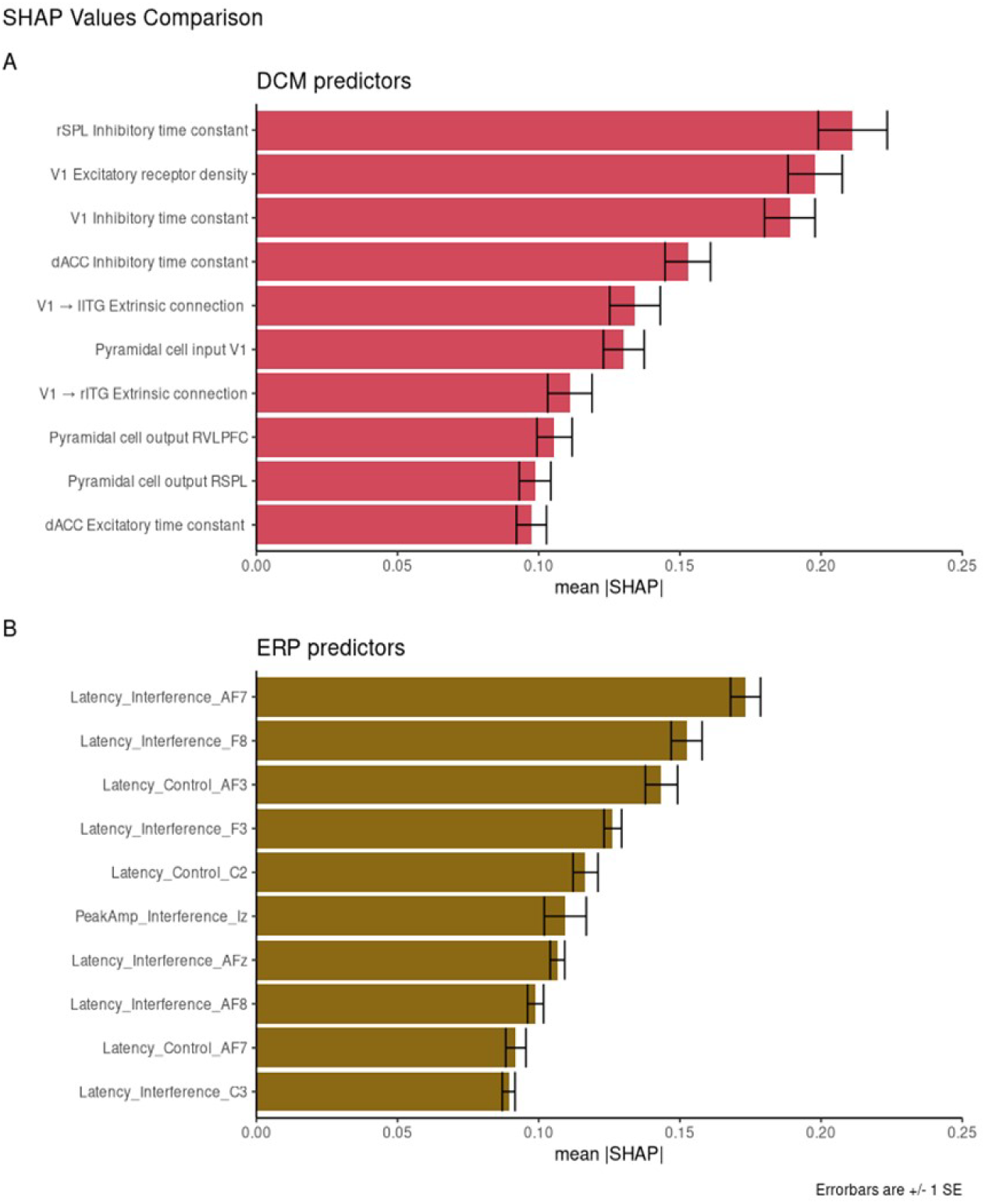
Feature Importance. SHAP values **for the** 10 best performing DCM features (A) and ERP features **(B).** Mean absolute SHAP values are reported with +/- 1 SE.

Individual DCM features have higher SHAP values that the EEG features with the same corresponding rank. For example, the most important feature of DCM predictors is the rSPL inhibitory time constant (mean SHAP = 0.22). The most important ERP predictor is AF7 Latency score in the interference condition (mean SHAP = 0.17). These results are directly comparable, with the larger SHAP score reflecting greater feature importance.

We also compared the mean SHAP scores between the two datasets, with DCM outperforming ERP across all ten most important features (the SHAP value of the top DCM feature is larger than the corresponding SHAP value for the top ERP feature; the SHAP value of the second-best DCM feature is larger than the corresponding SHAP value for the second-best ERP feature, etc.). We compared the mean absolute SHAP values in a paired t-test using rank as the pairwise grouping. The 10 top DCM features had higher SHAP values than the 10 top EEG features, as expected from a better predictive model (*t* = 4.28, *p* = 0.002, *CI =* [0.01, 0.03]).

Overall, DCM features appeared more powerful than EEG features (higher SHAP values). They also captured separate subtypes of depressed participants better. This may relate to DCM features’ greater ability to capture variability between individuals. Figure 5 shows the distributions of the ten most important DCM and EEG features. Visually, DCM features show distributions with central tendencies, with areas of non-overlap between patient and control distributions. This can be explained by the Laplace approximation used to define the posterior densities of DCM parameters.^77^ They occupy less of the available numeric range. ERP features are more uniformly dispersed over the available range.

**Figure 5:**
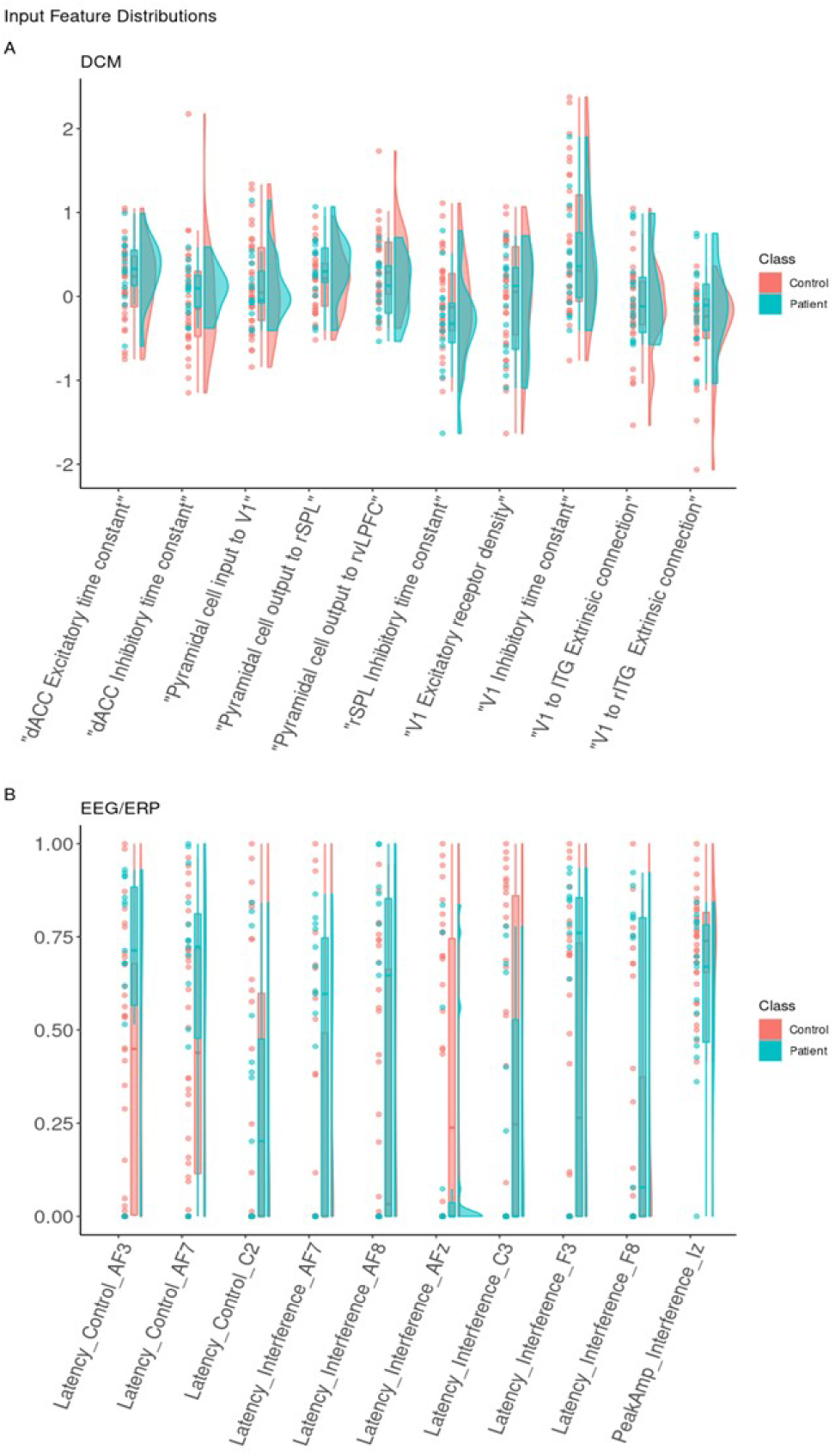
Feature distribution. Distributions for the 10 best DCM input features (A) and EEG/ERP input features (B). Classification features represented using dots for real observations alongside boxplots and density curves to show the shape of the distribution. Each is colored by class. Each input feature on the x-axis is shown with raw data points, boxplots, and density curves. Patients (blue) and controls (red) are shown separately. The EEG/ERP features are shown after min-max scaling to present latency and interference variables together on a comparable scale.

### Unsupervised Clustering

To capture depressive heterogeneity, *t-*SNE representations were made from unlabelled data of each of the 10 most important features for DCM and ERP feature sets. *T-*SNEs were generated using a perplexity value of 25 over 2,500 iterations. These embeddings were labelled post-hoc to determine if these features could elucidate differences between patients and controls. Figure 6 shows the three dimensional representations of patient embeddings in blue and control embeddings in red. Supplementary Figure 10 shows the recalculated t-SNE values for two dimensional representation. DCM feature embeddings show clustering tendencies, while EEG feature embeddings were more dispersed throughout the lower dimensional spaces with no clear patterns.

**Figure 6.**
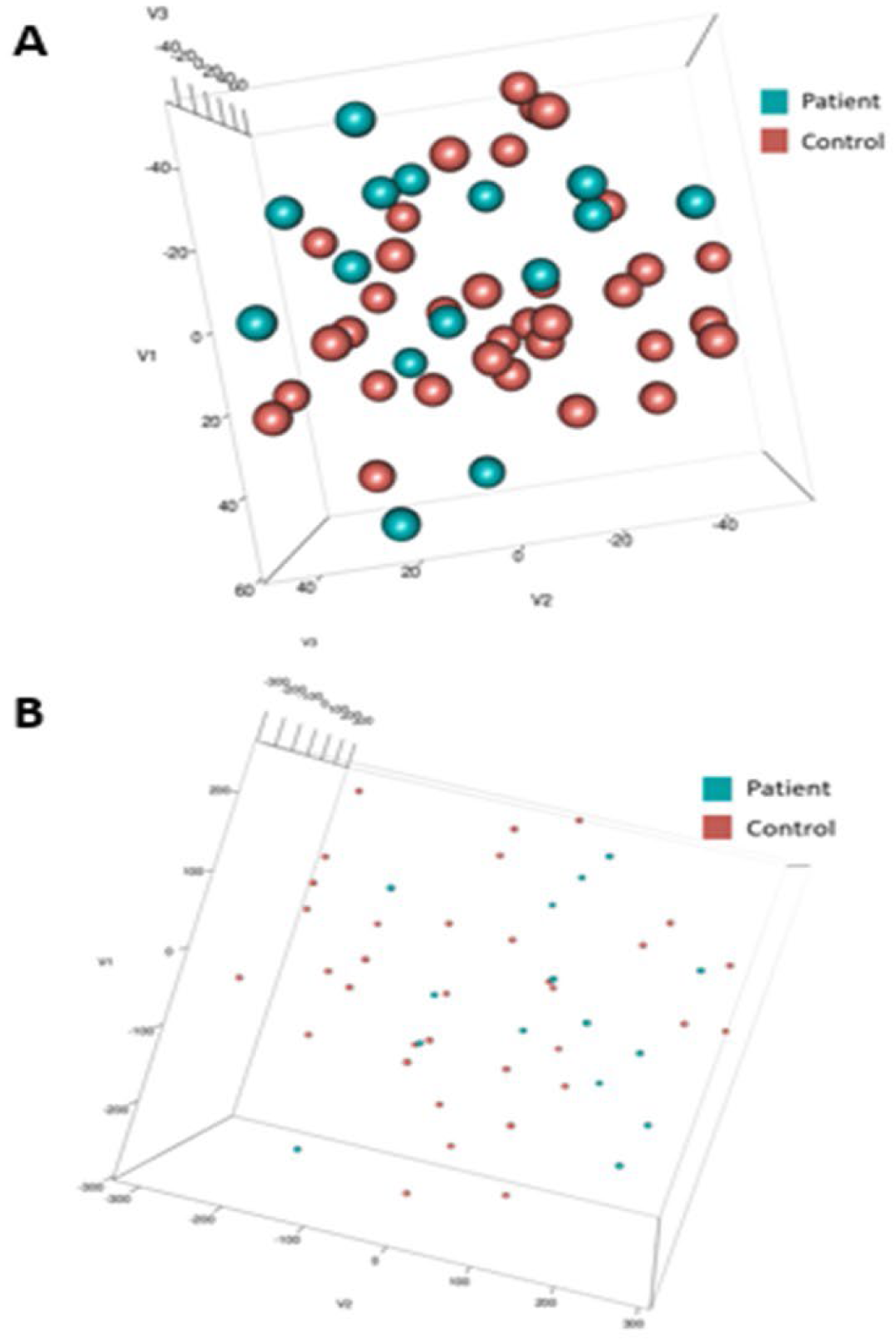
l-SNE embeddings. Three dimensional 1-SNE values for top 10 input features of DCM **(A)** and EEG/ERP **(B)** Post-hoc. dala points were colored blue (Patient) and red (Control). Note the difference in point size reflects the difference in axis scale (-/+ 40 in DCM compared with -/+ 300 in EEG).

This clustering pattern was specific to patients. We clustered the t-SNE embeddings using k-means and compared Silhouette scores for all participants and for the patient cohort only. Figure 7 shows the mean Silhouette score with +/- 1 standard error. Scores for both DCM and EEG results across 2-12 clusters (*k*) can be found in Suppl. Table 4. Silhouette scores decrease monotonically, suggesting that a two cluster representation is the most parsimonious solution in this small dataset. The lack of change in Silhouette scores over increasing values of *k* for the all-participants dataset suggests that there is no clear clustering solution that separates, e.g., controls and two or more patient types. This may reflect unstructured heterogeneity in the control participants.

**Figure 7:**
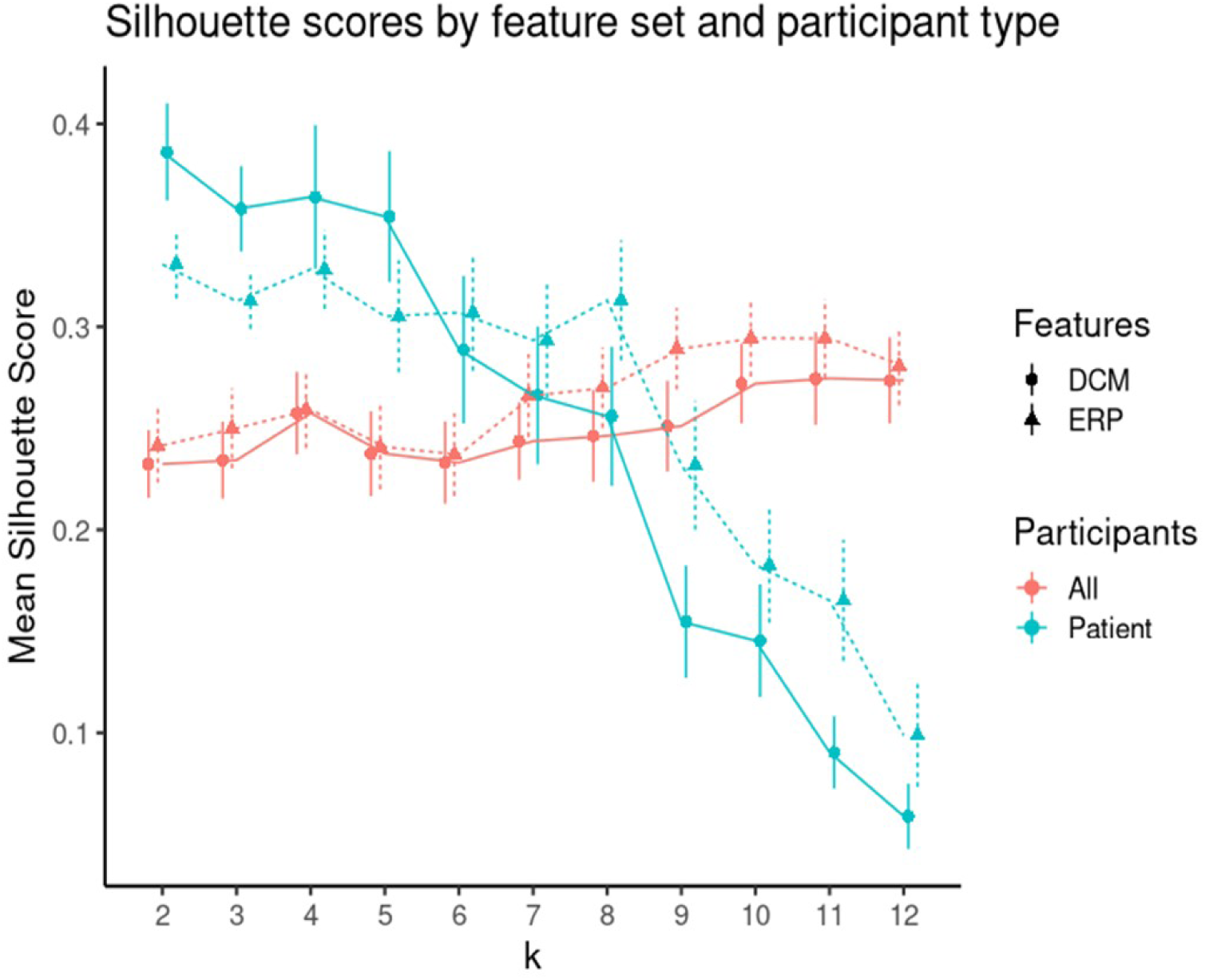
K-means clustering. Clustering was performed using varying numbers of *k* (from 2 to 12) to cluster 1) all participants, 2) only patients. To fmd the optimal k, we used Silhouette scores. Silhouette scores quantify the ratio of within over between cluster distance. A positive value signifies appropriate clustering [74], This figure shows the mean Silhouette scores (vertical axis) for each level of k (shown on tlie horizontal axis). Red dots show scores for patients and controls (all), while green dots show scores for patients only. Solid lines and discs depict scores for DCM features, while dashed lines and triangles for EEG features. Error bars show +/- 1 SE.

The DCM embeddings for the patients only at low *k* values had the highest mean Silhouette scores (*µ =* 0.388, *CI =* [0.334, 0.441]). Compared to the second highest Silhouette score, ERP embeddings for patients only, the DCM embeddings were significantly higher when compared using a two-sided t-test (*t =* 2.17, *p* = 0.032).

## Discussion

We demonstrated a proof of concept that transforming electrophysiological data to underlying biophysical parameters using DCM can more reliably capture variability that correlates with clinical status. Although our dataset is small and heterogeneous, DCM features outperformed raw EEG features on most classification metrics, and this held true across multiple classifier algorithms. Our results align with prior work that has successfully used DCM to identify differences between unipolar and bipolar depression.^22^ We considered participant-specific (first-level) analysis and group effects (second-level analysis) as two separate steps. An alternative approach could be to combine these steps into a single hierarchical model.^36, 77^ We will consider this elsewhere. Similarly, we used a Fixed Effects approach that could be replaced by Mixed-Effects.^7, 8^

If our DCM-based approach can be replicated on a larger dataset, it will provide an intriguing avenue for personalized medicine. There is a growing set of TMS and similar tools for manipulating brain connectivity.^79, 80^ For any individual patient, the approach we describe permits the identification of which DCM feature(s) are driving that patient’s vulnerability. Although this is still a speculative claim, it may be possible to then target and normalize those specific features. The feasibility of that approach will depend on whether these features change as a patient undergoes treatment or remain present even during euthymia.^81, 82^

We found that MSIT-induced variance was explained by feedforward connectivity from primary sensory and object discrimination areas to prefrontal cortex, plus changes in intrinsic, within-region connectivity. This is consistent with current working models of cognitive control, the construct tested by MSIT. In those models, information about cognitive control demands is computed posteriorly, then fed forward to anterior structures (dlPFC which then influences motor circuits).^83, 84^ Given that we explored a wide range of connectivity changes, our data-driven recovery of a known phenomenon provides some confidence that we identified known effects. At the same time, our analyses were based on prior assumptions about anatomically plausible connections. In future work, we will test these assumptions using BMS.

DCM suggested a low dimensional space of potential biomarkers (we went from 240 EEG-based to 92 biophysical features). This helps fitting prediction models on limited datasets. As an alternative to PCA^81, 85^, DCM-derived parameters are also directly interpretable; e.g. DCM synaptic connectivity or intrinsic excitability can directly map to potential treatments.^20^ Another DCM advantage is its biophysical models provide ways to test potential explanations of pathophysiology. This advantage can also be a limitation: for example, some prior knowledge about cortical regions and their interactions is required.

We emphasize that the classification performance of our models should not be treated as a claim that the current pipeline carries diagnostic or clinical utility. The sample size is too small; we and others have pointed out that small-sample-derived biomarkers do not generalize.^14, 86, 87^ Although we did perform some internal cross-validation, this sample size did not permit us to follow best-practices for preventing data leakage.^71^ Specifically, we performed SMOTE up-sampling on the full dataset, before conducting a training/validation split. On the other hand, the goal of this work was not to develop a reliable classifier and report its performance. The classification approach was used solely to compare the *relative* performance of DCM vs. ERP features (similar susceptibility to data leakage and effect size inflation).

Our feature importance scoring (SHAP) results should also be interpreted with caution. With this caveat, over half the highly weighted features from the DCM analysis came from more caudal regions and included within region features. This aligns with recent results in a larger dataset, where signals in primary sensory regions were more able to classify (non)response to antidepressant treatment than were signals from higher-order cognitive/associative regions.^88^ On the other hand, those response predictors were generally unstable in a cross-validation analysis.^89^ In all, we do not make claims that our specific identified markers/clusters are new generalizable findings. Rather, they are proof of concept for a larger point: that DCM parameters can be scored and interpreted, and that subtypes might be identifiable by clustering. With a larger dataset, it would become feasible to identify robust DCM features.^71^ There are such recent datasets available^90, 91^, particularly in depression^92, 93^. In future work we will consider these new datasets together with additional covariates, like stressful life events^94^, biological sex^6^, and others. These can easily be combined with the DCM pipeline we have shown here.

In summary, we have demonstrated the first proof of concept for a novel approach to identifying psychiatric biomarkers from EEG, based on converting manifest EEG signals to interpretable biophysical parameters. We demonstrated the potential superiority of this approach over the same biomarker pipeline applied to manifest data (in this case, ERPs). If applied to larger datasets and a more robust variety of data sources, this DCM-based pipeline can be an important new approach to dissecting the heterogeneity of depression and depressive vulnerability.

## Abbreviations

DCM: Dynamic Causal Modelling
ERP: Event-related Potentials
MSIT: Multi-Source Interference Task
JR: Jansen and Rit mass model
ICA: Independent Component Analysis
MCC: Matthew’s Correlation Coefficient
SHAP: Shapley Additive Explanation
PCA: Principal Component Analysis

## Data availability

This is a secondary analysis of a cohort collected in a previous study.^28^ No additional human participants review was required for this secondary analysis, as the primary study’s data had been deposited in an online repository (https://transformdbs.partners.org/), from which we obtained de-identified data. All participants gave informed consent for the primary study, which was overseen by the Massachusetts General Hospital Institutional Review Board.

## Supplementary Figures

**Supplementary Figure 1.**
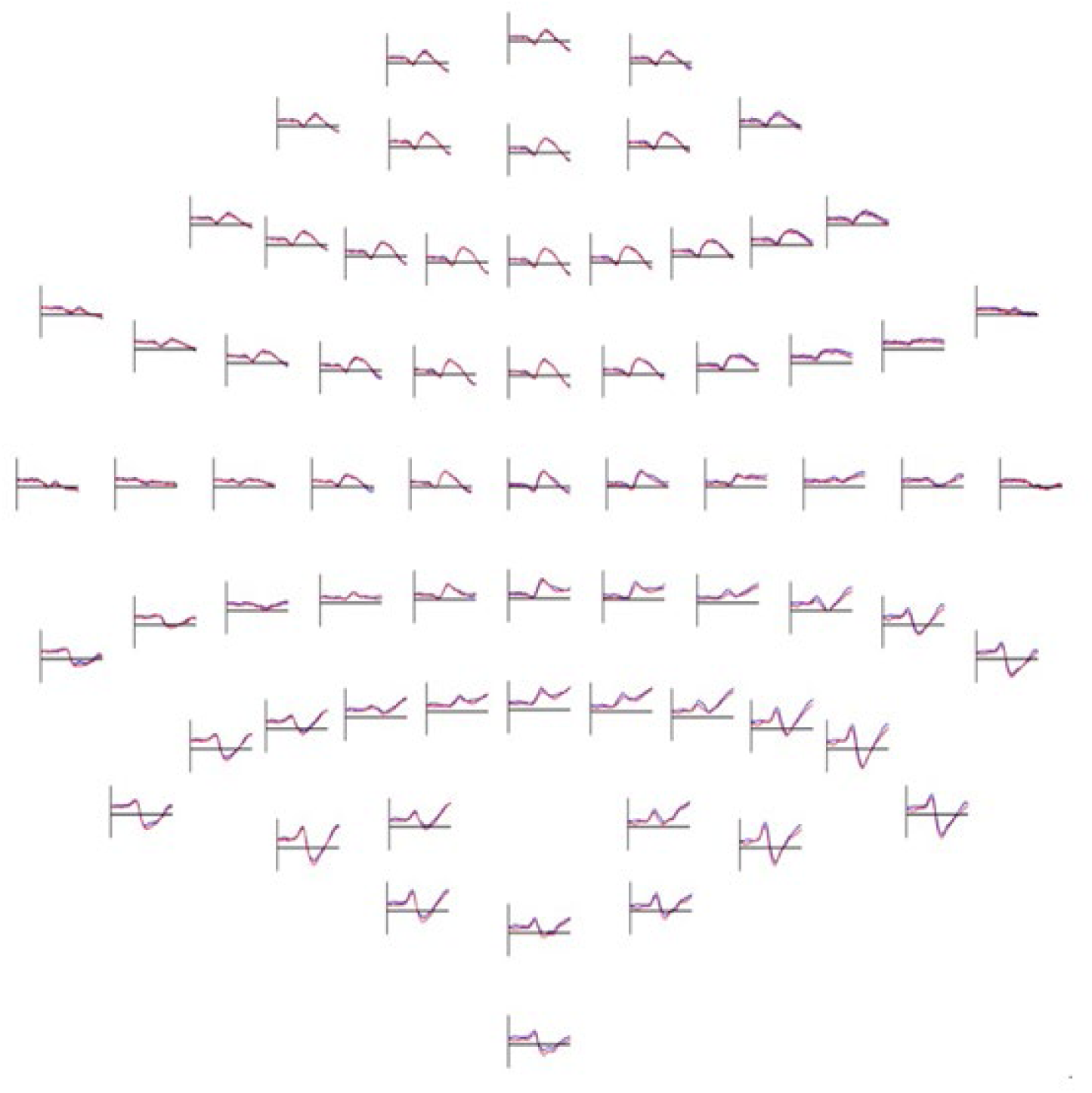
Topoplot of Mean ERP. EEG responses as topoplot for both control (blue) and interference condition (red) over 350 ms response time post stimulus onset.

**Supplementary Figure 2.**
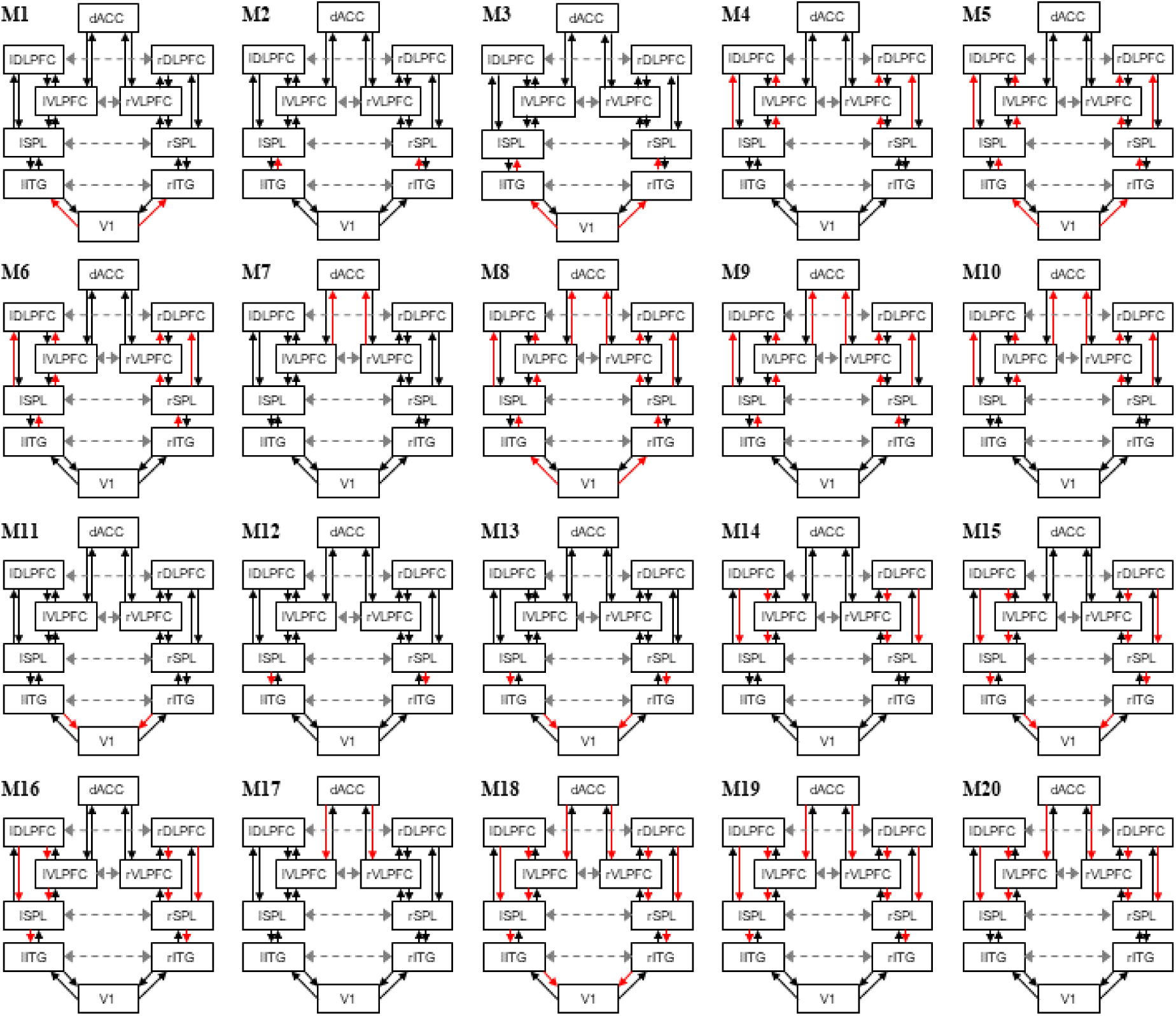
Example model space investigating extrinsic modulated connections. Candidate DCM models showing extrinsic modulated connections. They are variants of the functional network activated during the MSIT task. There were 12 extrinsic connections that could change between conditions. We assumed that these could change between one or more neighbouring pairs of brain areas: occipital to parietal, parietal to frontal and frontal to dACC. Connections that changed in each model are shown in red. See main text for explanation.

**Supplementary Figure 3.**
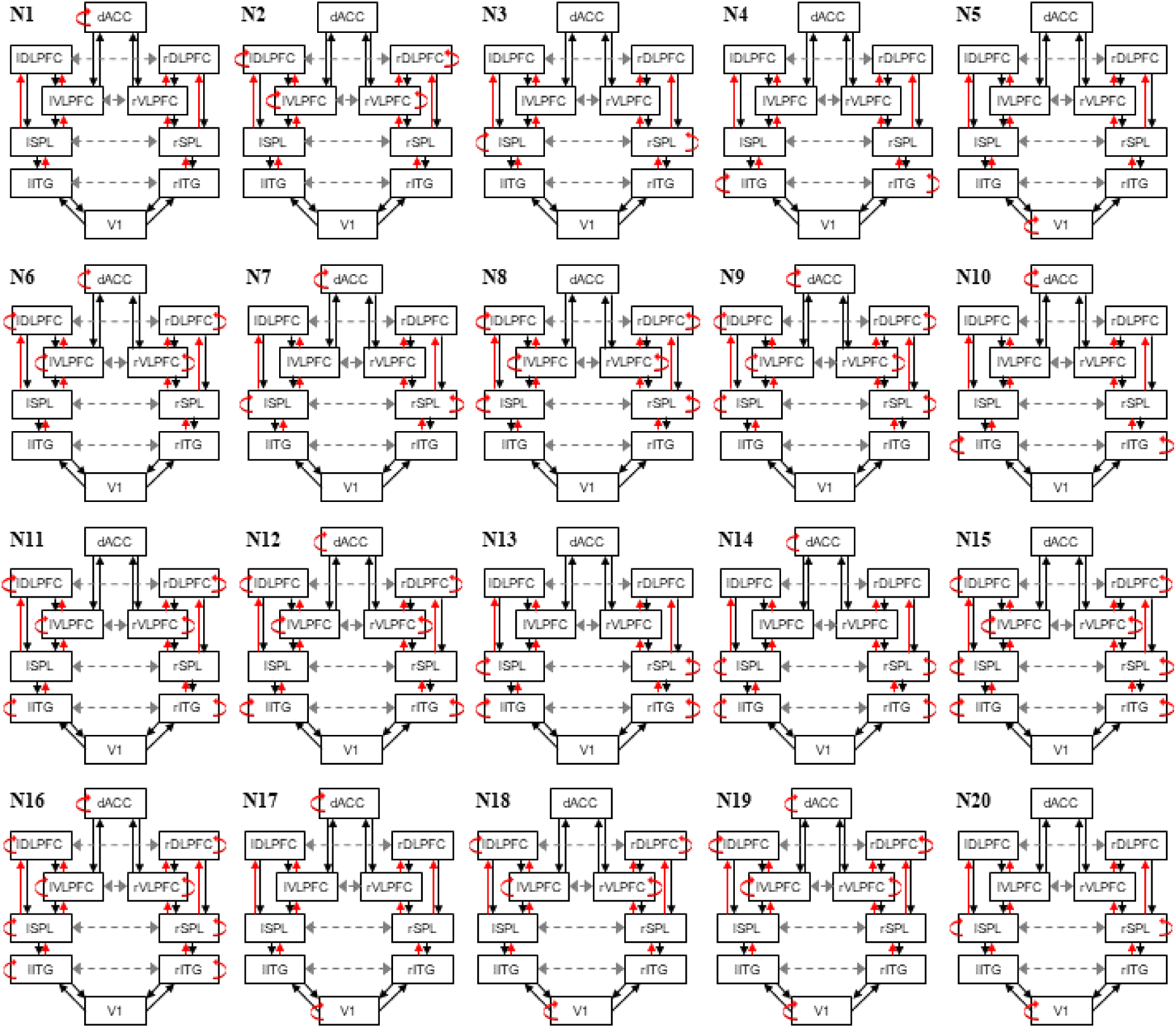
Example model space investigating intrinsic modulated connections. Candidate DCM models showing intrinsic modulated connections. Connections that could change are depicted in red, similarly to Supplementary Figure 2.

**Supplementary Figure 4.**
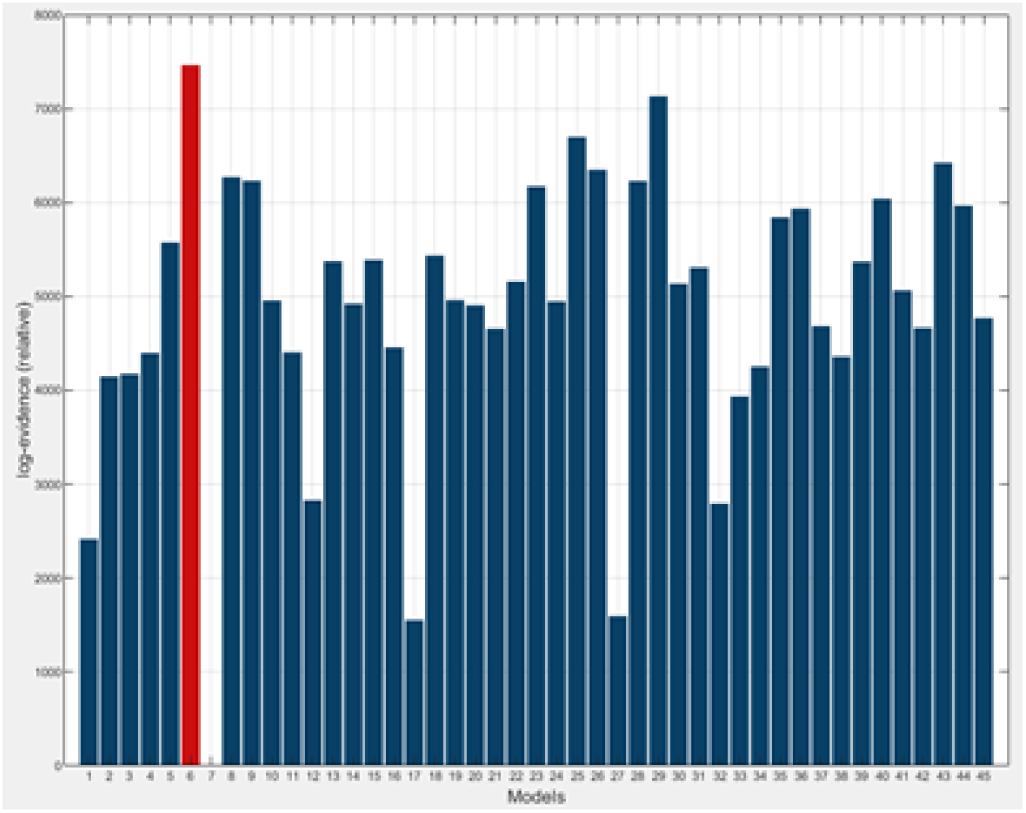
Extrinsic model selection (Controls). Bayesian model comparison among all models showing modulations in extrinsic connections. Model M6 is the model with the highest evidence (cf. Figure 2 in main text).

**Supplementary Figure 5.**
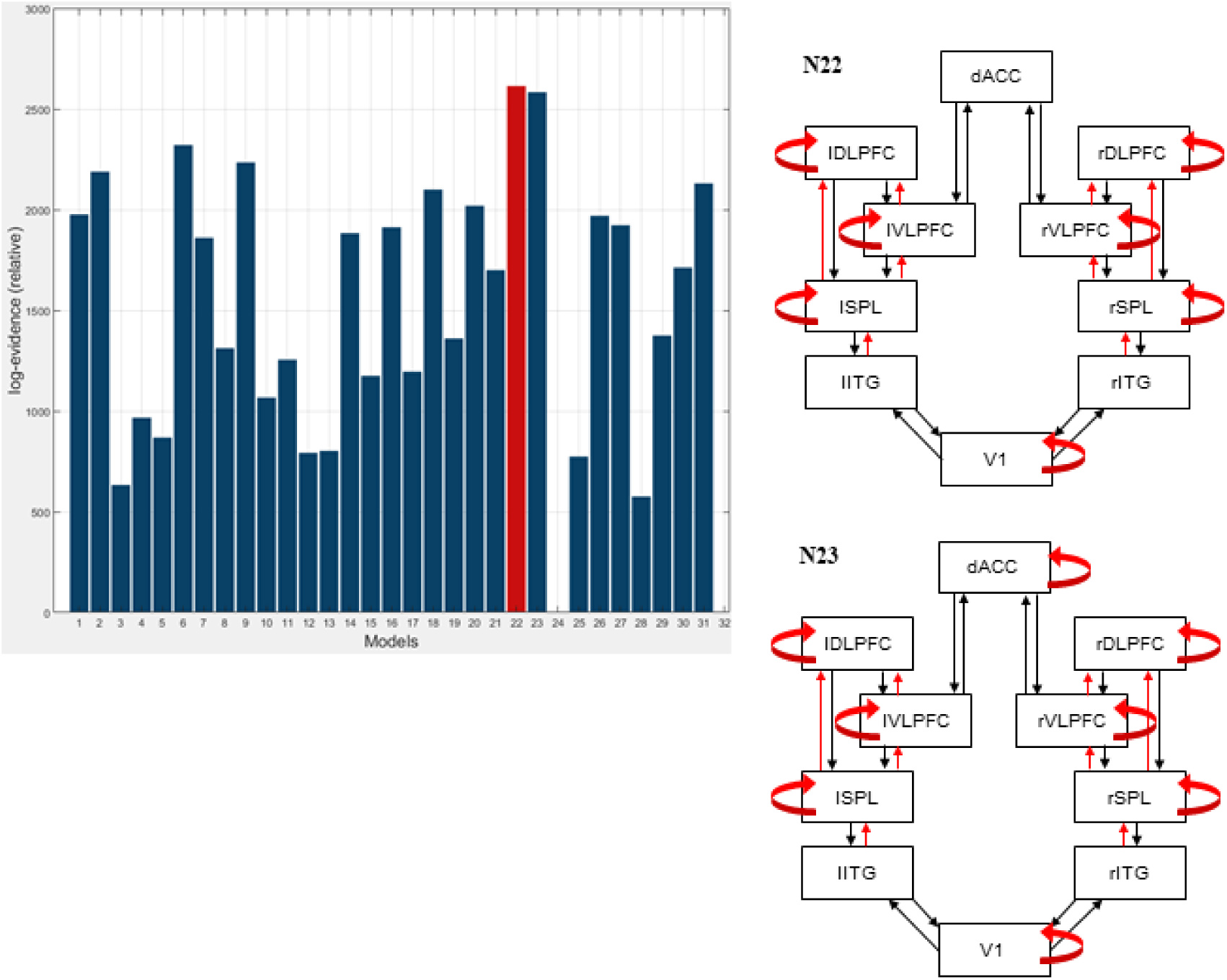
Intrinsic model selection (Controls). (Left) Each panel contains exemplar model predictions of neural activity produced by each population for the congruent (blue) and incongruent (red) conditions. Solid lines correspond to pyramidal cell predictions. Thick dashed to spiny stellates cells. The last pair of dashed lines corresponds to predictions from inhibitory interneurons.

**Supplementary Figure 6.**
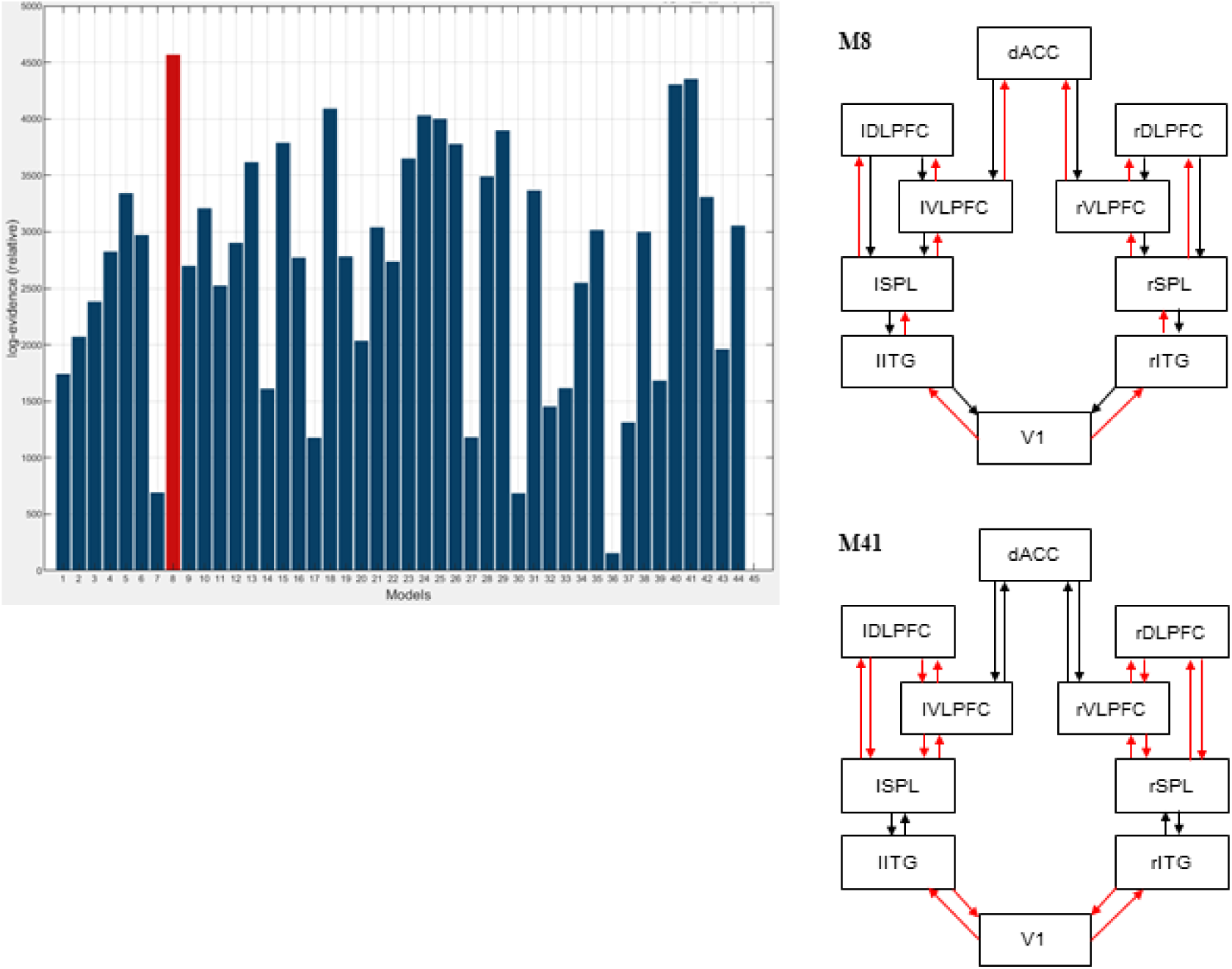
Extrinsic model selection (Patients) (Left) Similar to Suppl. Figure 4 for the Patient group. Model M8 has the highest evidence among all models showing modulations in extrinsic connections. (Right) Models M8 and its runner up M41.

**Supplementary Figure 7.**
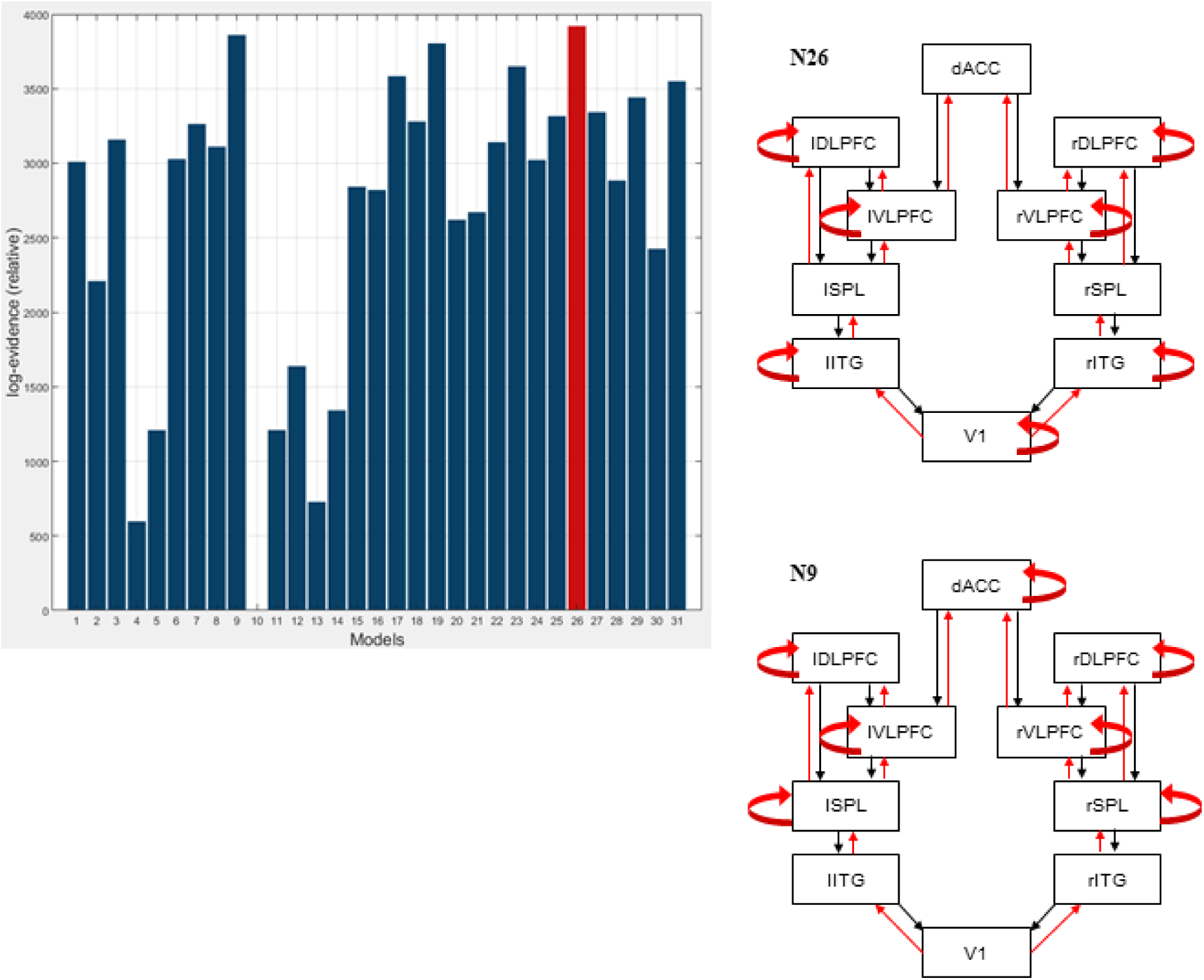
Intrinsic model selection (Patients) (Left) Similar to Suppl. Figure 5 for the Patient group. Model N26 has the highest evidence among all models showing modulations in intrinsic connections. (Right) Models N26 and its runner up N9.

**Supplementary Figure 8.**
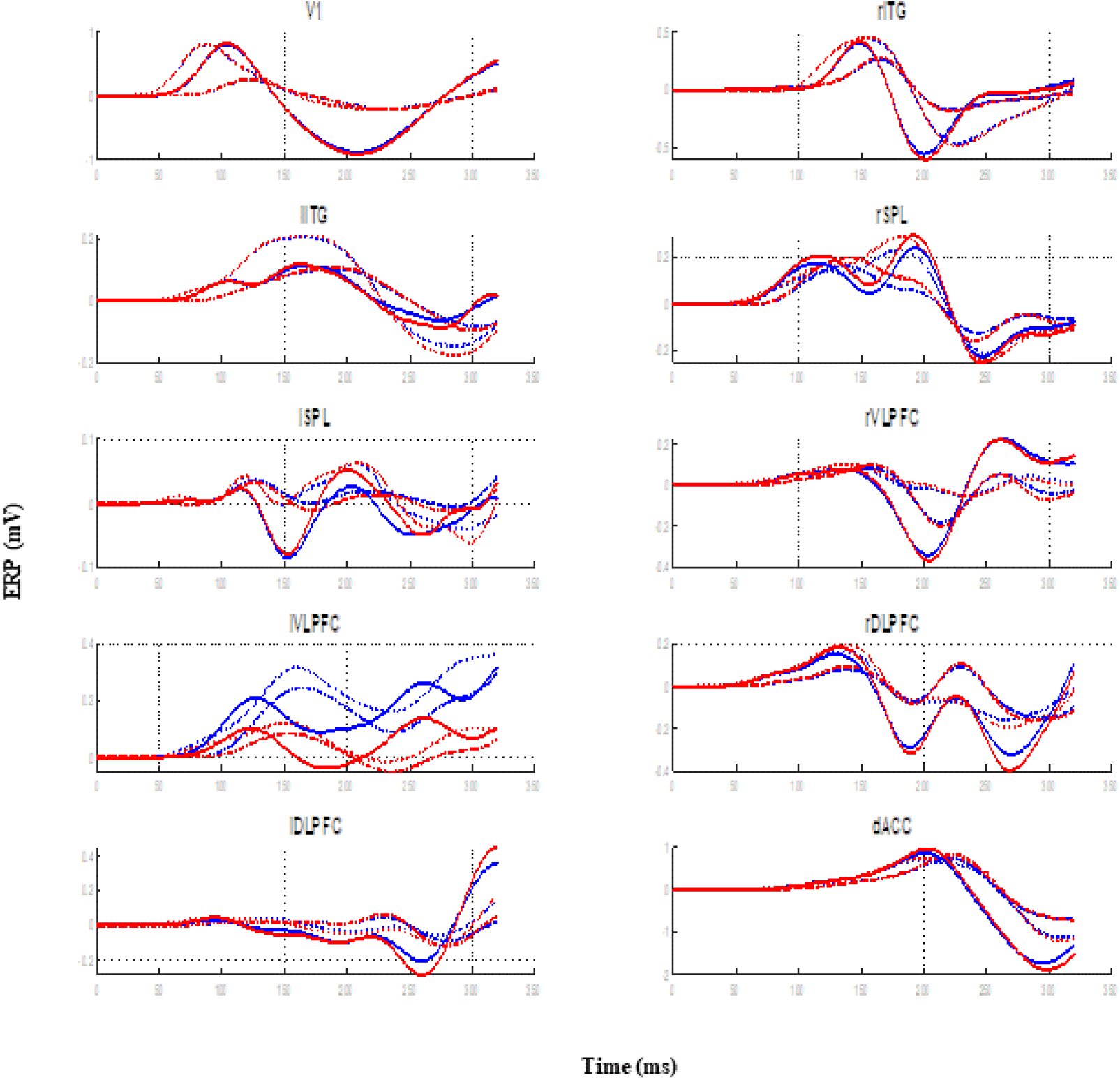
Each panel contains exemplar model predictions of neural activity produced by each population for the congruent (blue) and incongruent (red) conditions. Solid lines correspond to pyramidal cell predictions. Thick dashed to spiny stellates cells. The last pair of dashed lines corresponds to predictions from inhibitory interneurons.

**Supplementary Figure 9.**
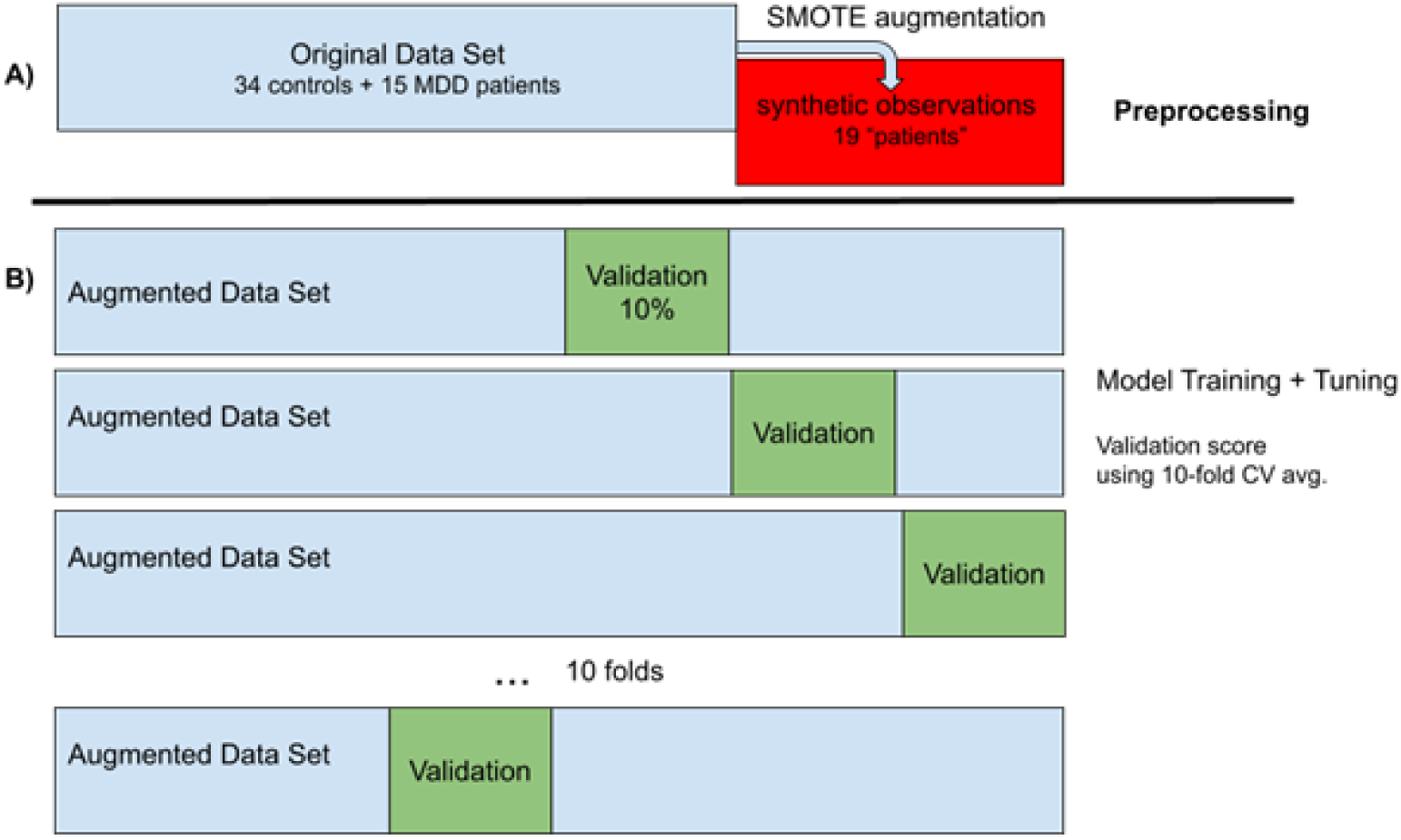
**Data Sampling Strategy**. Data segmentation plan and model training design. A) The original data set of 34 controls and 15 MDD patients was augmented using SMOTE to create 19 additional synthetic participants during preprocessing. B) The augmented data was split into 10 separate cross-validation folds by withholding a different, random, 10% of the data for performance measurement and using the remaining 90% for model training and hyperparameter tuning.

**Supplementary Figure 10.**
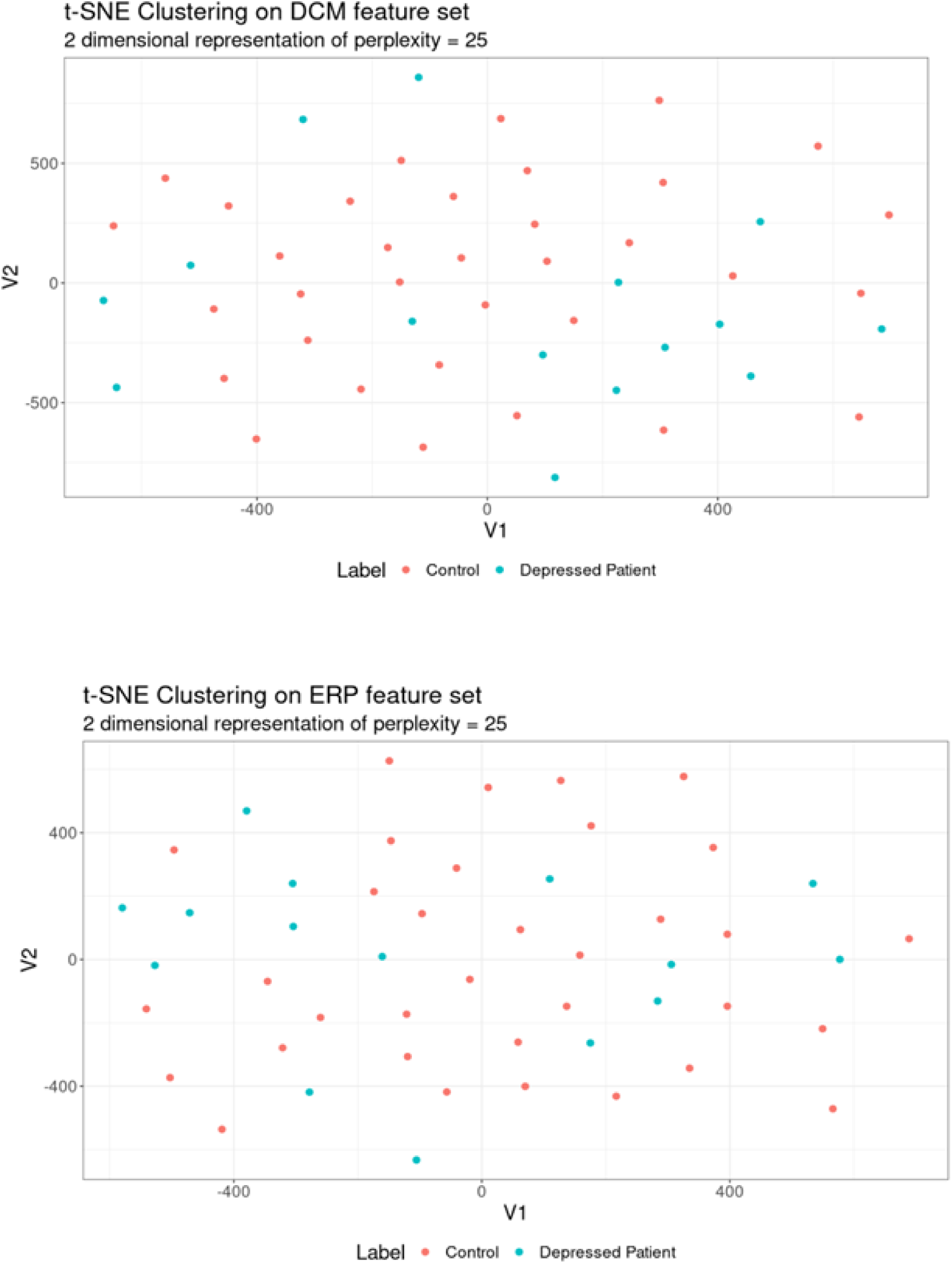
Two dimensional t-SNE representations. Two dimensional t-SNE values for top 10 input features of DCM (top) and EEG/ERP (bottom). Post-hoc, data points were colored blue (MDD patients) and red (Control)

---

**Supplementary Table 1.** Wilcoxon-tests for differences in ERF latencies between control and interference conditions.

| <u>Channel</u> | <u>Control – Interference Latency (s)</u> | <u>statistic</u> | <u>p-value</u> | <u>CI low</u> | <u>CI high</u> |
| --- | --- | --- | --- | --- | --- |
| P3 | 0.027 | 347 | 0 | 0.014 | 0.041 |
| C3 | 0.032 | 830 | 0 | 0.016 | 0.043 |
| FCz | 0.036 | 663 | 0 | 0.023 | 0.048 |
| F1 | 0.037 | 724 | 0 | 0.022 | 0.051 |
| FT7 | 0.031 | 752 | 0 | 0.018 | 0.044 |
| Fz | 0.033 | 637 | 0 | 0.018 | 0.048 |
| F7 | 0.026 | 744 | 0 | 0.011 | 0.042 |
| FC3 | 0.026 | 806 | 0 | 0.008 | 0.04 |
| C1 | 0.039 | 620 | 0 | 0.019 | 0.057 |
| C5 | 0.031 | 698 | 0.001 | 0.013 | 0.043 |
| CP3 | 0.017 | 422 | 0.001 | 0.008 | 0.033 |

**Supplementary Table 2.** **EEG Channels Retained for Classification** AF3 C2 AF7 FTP P03 F8 02 AFz F4 AES C4 Oz PS Iz PO7 TP9 F3 Pl TP10 F7 ET8 C3 Fp2 P5

**Supplementary Table 3.**
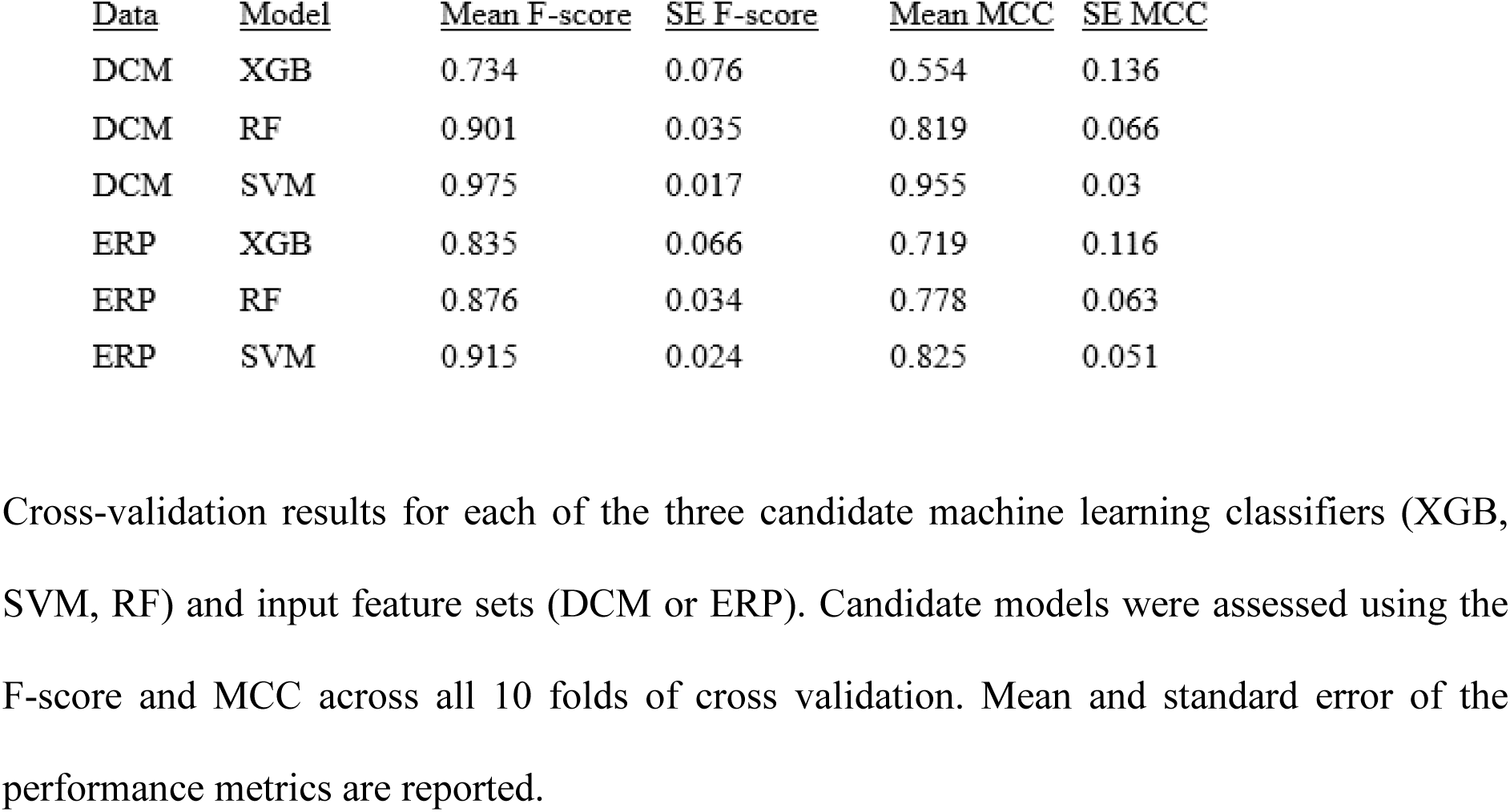
Cross-validation Scores

**Supplementary Table 4.** Silhouette Scores

| **<u>Participants</u>** | **<u>Features</u>** | **<u>k</u>** | **<u>Mean Silhouette Score</u>** | **<u>SD</u>** | **<u>SE</u>** | **<u>CI min</u>** | **<u>CI max</u>** |
| --- | --- | --- | --- | --- | --- | --- | --- |
| All | DCM | 2 | 0.244 | 0.111 | 0.016 | 0.276 | 0.213 |
| All | DCM | 3 | 0.226 | 0.136 | 0.020 | 0.264 | 0.187 |
| All | DCM | 4 | 0.256 | 0.138 | 0.020 | 0.295 | 0.217 |
| All | DCM | 5 | 0.240 | 0.135 | 0.019 | 0.278 | 0.202 |
| All | DCM | 6 | 0.236 | 0.140 | 0.020 | 0.275 | 0.196 |
| All | DCM | 7 | 0.262 | 0.135 | 0.019 | 0.300 | 0.224 |
| All | DCM | 8 | 0.242 | 0.147 | 0.021 | 0.284 | 0.201 |
| All | DCM | 9 | 0.260 | 0.150 | 0.022 | 0.302 | 0.218 |
| All | DCM | 10 | 0.279 | 0.125 | 0.018 | 0.314 | 0.243 |
| All | DCM | 11 | 0.276 | 0.140 | 0.020 | 0.315 | 0.236 |
| All | DCM | 12 | 0.296 | 0.137 | 0.020 | 0.335 | 0.258 |
| Patient | DCM | 2 | 0.388 | 0.189 | 0.027 | 0.441 | 0.334 |
| Patient | DCM | 3 | 0.371 | 0.194 | 0.028 | 0.426 | 0.316 |
| Patient | DCM | 4 | 0.360 | 0.180 | 0.026 | 0.411 | 0.309 |
| Patient | DCM | 5 | 0.335 | 0.219 | 0.032 | 0.397 | 0.273 |
| Patient | DCM | 6 | 0.285 | 0.183 | 0.026 | 0.337 | 0.233 |
| Patient | DCM | 7 | 0.274 | 0.189 | 0.027 | 0.327 | 0.220 |
| Patient | DCM | 8 | 0.235 | 0.212 | 0.031 | 0.295 | 0.174 |
| Patient | DCM | 9 | 0.162 | 0.196 | 0.028 | 0.217 | 0.106 |
| Patient | DCM | 10 | 0.146 | 0.187 | 0.027 | 0.198 | 0.093 |
| Patient | DCM | 11 | 0.096 | 0.151 | 0.022 | 0.138 | 0.053 |
| Patient | DCM | 12 | 0.050 | 0.092 | 0.013 | 0.076 | 0.023 |

| <u>Participants</u> | <u>Features</u> | <u>k</u> | <u>Mean Silhouette Score</u> | <u>SD</u> | <u>SE</u> | <u>CI min</u> | <u>CI max</u> |
| --- | --- | --- | --- | --- | --- | --- | --- |
| All | ERP | 2 | 0.245 | 0.131 | 0.019 | 0.282 | 0.208 |
| All | ERP | 3 | 0.253 | 0.144 | 0.021 | 0.294 | 0.213 |
| All | ERP | 4 | 0.243 | 0.139 | 0.020 | 0.282 | 0.204 |
| All | ERP | 5 | 0.246 | 0.136 | 0.020 | 0.285 | 0.208 |
| All | ERP | 6 | 0.256 | 0.141 | 0.020 | 0.296 | 0.216 |
| All | ERP | 7 | 0.262 | 0.139 | 0.020 | 0.301 | 0.222 |
| All | ERP | 8 | 0.263 | 0.119 | 0.017 | 0.297 | 0.230 |
| All | ERP | 9 | 0.247 | 0.148 | 0.021 | 0.289 | 0.205 |
| All | ERP | 10 | 0.256 | 0.158 | 0.023 | 0.300 | 0.211 |
| All | ERP | 11 | 0.272 | 0.148 | 0.021 | 0.313 | 0.230 |
| All | ERP | 12 | 0.275 | 0.145 | 0.021 | 0.316 | 0.234 |
| Patient | ERP | 2 | 0.322 | 0.094 | 0.014 | 0.349 | 0.296 |
| Patient | ERP | 3 | 0.288 | 0.121 | 0.017 | 0.323 | 0.254 |
| Patient | ERP | 4 | 0.325 | 0.114 | 0.016 | 0.357 | 0.293 |
| Patient | ERP | 5 | 0.346 | 0.157 | 0.023 | 0.390 | 0.302 |
| Patient | ERP | 6 | 0.353 | 0.171 | 0.025 | 0.401 | 0.305 |
| Patient | ERP | 7 | 0.306 | 0.213 | 0.031 | 0.366 | 0.245 |
| Patient | ERP | 8 | 0.294 | 0.220 | 0.032 | 0.356 | 0.232 |
| Patient | ERP | 9 | 0.266 | 0.229 | 0.033 | 0.331 | 0.201 |
| Patient | ERP | 10 | 0.209 | 0.204 | 0.029 | 0.267 | 0.152 |
| Patient | ERP | 11 | 0.175 | 0.216 | 0.031 | 0.236 | 0.114 |
| Patient | ERP | 12 | 0.105 | 0.180 | 0.026 | 0.156 | 0.054 |

Mean Silhouette scores for the EEG/ERP clusters across the varying number of clusters (k) for all participants and only the patient cohort. Standard deviation, standard error and 95% confidence interval min and max are also reported.

## Supplementary Methods

### Dataset and Task

The dataset included was generated from 15 psychiatric patients who reported current or past depressive symptoms and 34 non-diagnosed controls. Importantly, this dataset was not limited to patients diagnosed with unipolar depression, but included bipolar and unspecified depression. We considered this a better demonstration of our approach to heterogeneity. This is a secondary analysis of a cohort collected in a previous study^28^. No additional human participants review was required for this secondary analysis, as the primary study’s data had been deposited in an online repository (https://transformdbs.partners.org/), from which we obtained de-identified data. All participants gave informed consent for the primary study, which was overseen by the Massachusetts General Hospital Institutional Review Board.

EEGs were recorded with a 70-channel electrode cap, based on based on the 10–10 electrode-placement system (Easycap, Vectorview, Elekta-Neuromag, Helsinki, Finland), using a sampling rate of 1000 Hz. All recordings were completed in an electro-magnetically shielded room, with individual channel impedances kept below 5000 Ohm.

EEGs were collected as participants performed the Multi-Source Interference Task (MSIT).^29^ The task requires participants to report on a presented stimulus by using their index, middle or ring finger to press three buttons corresponding to numbers 1, 2, and 3 respectively. The stimulus appears on a screen displaying three numbers; one number (the target) is different from the other two (distractors). The participant identifies the target by pressing the corresponding buttons. There are two task conditions, control and interference, which are interleaved to prevent development of response sets. During control trials, the distractor numbers are zeros and the location of the target number is aligned with its corresponding button. In interference trials, the distractors are non-zero numbers (valid targets; flanker effect), and the target is in a location misaligned with that of the button (Simon effect). The dataset was provided in a pre-processed state whereby artefacts had been removed using Independent Component Analysis (ICA) and bad trials (e.g., excessive rates of voltage change suggesting artefact) removed. The continuous signal data had been segmented into epochs using a time window of 1.5 seconds before stimulus onset and 2 seconds after. No further artefact rejection was performed in this study.

### ERP Analysis

P300 components were extracted from 70 EEG channels (average ERPs over participants are included in Figure 1A; see also Suppl. Figure 1). Previous MSIT studies found differences in the P300 component between task conditions.^32, 33^ P300s are a common signature of conflict and cognitive control, and are believed to arise from mid-frontal processing of incongruent stimuli.^34, 35^ The P300 signal was defined as evoked responses between 250ms - 350ms after event onset. Only trials with correct responses were used. Wilcoxon tests for the effect of conditions (interference vs. control), on each channel across controls and patients showed that latency (but not peak amplitude) exhibited significant differences, see Suppl. Table 1. *P*-values were corrected for multiple comparisons using a Bonferroni corrected threshold value of *p<*0.0007.

### Dynamic Causal Modelling (DCM)

We used Dynamic Causal Modelling (DCM)^20–22, 36–41^, which allows one to infer processes at the neuronal level from scalp EEG measurements^2^. DCM models the changes of intrinsic (within area) and extrinsic (between area) connections across task conditions. It allows one to assess whether information flow changes in the same way (top-down, bottom-up or both) between the two task conditions across all participants. DCM includes a biophysical model that predicts the neuronal activity that underlies the observed EEG signal.^42^ This model comprises a Jansen and Rit (JR) neural mass that included three populations of neurons: excitatory pyramidal cells, excitatory spiny stellate cells, and inhibitory interneurons (smooth stellate cells). These populations are connected with one another for excitatory (black) and inhibitory (red) connections, and also with populations in other areas (Figure 1B). JR models can predict both evoked and induced responses and have been used in theoretical and experimental studies.^27, 43–46^ DCM was implemented using SPM12.

Beyond DCM, the JR model has been used to describe complex brain dynamics including epilepsy^47^ and TMS effects.^48^ It was the first model of a cortical circuit that produced a variety of brain dynamics (evoked responses, attractor states, spike-wave discharges etc.) by simply varying its parameters. In the context of DCM, the JR model has been used to predict ERPs during attention and other cognitive tasks, including Mismatch Negativity (MMN).^25, 49, 50^

DCM also includes an observation model that transforms neuronal activity predictions from the biophysical model above to a predicted scalp-observed EEG signal. We here used the JR model and DCM to fit ERPs during the MSIT. Our analysis focused on time domain data. The observation model for each brain area corresponds to an equivalent current dipole (ECD).^51^ There are as many dipoles as brain areas. Each dipole has 6 parameters: 3 for its location, 2 for its orientation and 1 for each amplitude. For more details, see ^52^. By estimating the parameters of both models simultaneously, we can separate different sources of variability in the EEG signal like neuronal dynamics, volume conduction and other sources of noise observed with EEG. Fitting exploits a non-linear optimization (Bayesian) approach.^32^ Briefly, neuronal dynamics are prescribed by the JR model. Then they are projected to the EEG sensor level via an ECD model. Both models (JR and ECD) are included in DCM. Their parameters are estimated simultaneously to account for conditional dependencies. Because DCM estimates both the observation and neural models, it effectively performs source reconstruction from EEG electrodes to neural sources. Their parameters are estimated simultaneously to account for conditional dependencies.

DCM was implemented using SPM12. The functional network modelled with DCM can be seen in Figure 2 (cf. model M1 in top left corner, all other models include the same network and assume changes in different connections, explained below). This network is comprised of areas activated during the MSIT.^29, 53^ Changes in functional connectivity within this network were observed, at the group level, in patients with depression.^42, 54–56^ The following anatomical regions were selected (in MNI coordinates)^35^: primary visual cortex (V1) [24, -83, 7], right inferior temporal gyrus (rITG) [52, -54, -14], left inferior temporal gyrus (lITG) [- 54, -33, -25], right superior parietal lobule (rSPL) [23, -59, 56], left superior parietal lobule (lSPL) [-33, -58, 57], right ventrolateral prefrontal cortex (rVLPFC) [44, 25, -12], left ventrolateral prefrontal cortex (lVLPFC) [-44, 35, -5], right dorsolateral prefrontal cortex (rDLPFC) [35, 41, 18], left dorsolateral prefrontal cortex (lDLPFC) [-28, 48, 4] and dorsal anterior cingulate cortex (dACC) [2, 3, 53]. We did not model other medial cortical structures (e.g., dorsomedial PFC or supplementary motor area) because it was not clear that their activity could be reliably disambiguated from underlying dACC. We similarly combined midline structures where left/right disambiguation would be uncertain. We considered a parsimonious brain network that included areas from each level of the cortical hierarchy – sensory, temporal, parietal, dorsal and ventral frontal areas, and ACC. Our assumption was that ERPs resulted from coordinated activity across this hierarchy. EEG recordings reveal their temporal profile, which, in turn, is the result of different temporal scales characterizing dynamics in each brain area. This affects signal propagation in different parts of the network and might be different in participants whose depressive symptoms arise from different network dysfunctions. Such differences, if they existed, can be revealed by DCM and the machine learning algorithms used below. This is because DCM estimates neural parameters after accounting for the observation model.

The biophysical model included in DCM predicts P300 responses. Data on the other hand, included ERP recordings from different participants, patients and controls. By fitting the DCM model predictions to this data, the variability in ERP recordings results in variability in the biophysical model parameter estimates across participants. That is, apparent noise or heterogeneity in the scalp-level recordings might arise from a small number of disruptions in the underlying network, which could be considered as biotypes or endophenotypes of depression.

Because we did not know how connectivity changed between task conditions, we compared several variants of the same biophysical model describing the network of Supplementary Figure 2. We considered a network containing all of our modelled brain regions: V1, ITG, SPL, vlPFC, dlPFC and dACC. We assumed forward and backward connections between specific areas, as well as lateral connections between homologous areas in the right and left hemispheres. The model variants differed in the connections that could vary between the two task conditions. Following Pinotsis, Buschmann, and Miller^40^, we first considered changes of extrinsic connections (i.e. between nodes) only. The first twenty candidate models we considered are shown in Supplementary Figure 2. There were 12 extrinsic connections that could change between conditions. We assumed that these could change between one or more neighbouring pairs of brain areas: occipital to parietal, parietal to frontal and frontal to dACC, that is, V1◊ ITG, ITG◊SPL, SPL ◊{vlPFC, dlPFC} and vlPFC◊ dACC. Connections that were permitted to change in each model are depicted with red arrows. Feedforward connections are depicted with bottom-up arrows, feedback connections with top-down arrows. Assuming that only the forward connections changed between MSIT conditions, we obtained models M1-M10 in Supplementary Figure 2. Only bottom-up arrows are red. Assuming that only backward conditions changed, we then obtained models M11-M20. Here, only top-down arrows are red. Similarly, assuming that both forward and backward connections changed, we obtained another ten models. These include ten models similar to those in Supplementary Figure 2, where both bottom-up and top-down arrows are red. The remaining fifteen models include models where we relaxed the constraint that between-condition differences would only be reflected in connections between neighbouring pairs of brain areas. in the first five models, we assumed that feedforward connections between V1→ ITG changed along with one of the five core connections, ITG◊SPL, SPL ◊vlPFC, SPL ◊dlLPFC, vlPFC◊ dlPFC and vlPFC◊ dACC. The next five models were similar, where instead of feedforward we considered feedback connections. The last five models included assumptions that both feedforward and feedback connections changed. Overall, the candidate model space comprised 45 models. To sum up, we compared all candidate models where forward or backward connections changed between different parts of the brain network. This yielded the extrinsic connections that were modulated during the task. After comparing these 45 models, we found one that could explain the data best (had the highest evidence). Then, we considered variations of this model assuming changes in intrinsic connections from each node to itself (in addition to changes in extrinsic connections that the winning model above assumed). We thus assumed that intrinsic connections could change at any (combination of) brain areas: V1, ITG, SPL, {vlPFC, dlPFC} and dACC. For example, assuming that intrinsic connections changed in only one of the above five brain areas (the two PFC subareas considered together), we obtained models N1-N5 in Supplementary Figure 3. The remaining models are just all possible combinations of the above five models. Thus, we compared 32 candidate models in total.

### Bayesian Model Selection (BMS)

For model comparison, we used an approach known as Bayesian model selection (BMS). This was performed assuming fixed-effects (FFX).^57^ BMS fits competing models to EEG data and assesses the most likely model. This exploits a Bayesian cost function (called relative Free Energy^33^), which quantifies model evidence (i.e., the Free Energy is a score of model fits). The Free Energy is similar to the Bayesian Information Criterion, BIC, in that it includes a complexity term dependent on both the number of parameters and their posterior correlations. That is, models with more free parameters are automatically penalized compared to models with fewer. Then, the most likely model (among a set of candidate models) is the one with the largest Free Energy. This is quantified in terms of the Bayes Factor (BF). This is a Bayesian analogue of the usual odds ratio and quantifies the probability that one out of many models has generated the EEG data. It provides the (relative) probability that this could be a true model of the brain compared to alternative models. Using BF, we can pool together evidence of the most likely model across different participants. BF is defined as the ratio of the Free Energy of two models. If BF*(1vs2)*>3, we can say that model 1 is better than 2—or more exactly, there is strong evidence for 1 relative to 2, see^40^. Note that the Free Energy can be thought of as an alternative to cross-validation in terms of preventing overfitting. Cross-validation assesses how well a model can explain unseen data, while Free Energy yields the most parsimonious model to explain observed data by penalizing for complexity.^58^ These two processes have been shown to be mathematically equivalent.^59^

### Feature Sets

Two distinct data sets of input features were used to train machine learning models, DCM parameters and EEG features. DCM parameters from individual subjects were obtained after fitting the biophysical model to EEG data, see previous section. Recall that the model included 12 extrinsic connections in each hemisphere connecting 10 brain areas in total. Each of these areas included a microcircuit like the one shown in Figure 1B. This includes 10 excitatory and inhibitory connections, each characterized by a synaptic time constant and synaptic efficacy, and recurrent connectivity parameters characterizing the gain of pyramidal cells and inhibitory interneurons. To sum up, the biophysical model included the following parameters identified for each participant: extrinsic connectivity, *A* (12x2=24 parameters), differences in extrinsic connectivity between MSIT conditions, *B* (24 parameters, derived from the model fitting as shown in Results), excitatory and inhibitory receptor density, *G* (2x10=20 parameters), strength of connections between the three populations of the JR model shown in Figure 1B, *H* (4 parameters; see arrows in Figure 1B), and excitatory and inhibitory synaptic time constants, *T* (20 parameters). The above parameters were used as predictor features in machine learning algorithms below. We thus obtained 92 DCM predictor features. The DCM features used included intrinsic and extrinsic connections that were found to differ between MSIT conditions in both the patient and control DCM fits.

To compare the predictive power of the DCM parameter estimates against the EEG features, we used an equal number of ERP features. The full set of potential EEG features included 240 variables (60 EEG Channels x 2 conditions x 2 variables, i.e. ERP peak amplitude and latency differences between the two MSIT conditions). We reduced the number of channels to 24 so that the total number of EEG features was similar to the number of DCM parameters to reduce bias when comparing ERP and DCM feature sets. We performed permutation testing to select the channels of greatest information gain.^60, 61^ The ERP features were chosen based on their contribution to a random forest model (constructed without hyperparameter tuning). This “naive” random forest allowed us to select channels with features that were most beneficial in separating classes while still allowing for multiple interaction effects between features. The tradeoff of this method stems from using a reduced number of features with the benefit that they are potentially more meaningful, and easier to interpret. The selected channels are included in Supplementary Table 2.

### SMOTE Oversampling and Model Training

Out of the 49 participants, only 15 were patients with depressive symptoms. The dataset was thus imbalanced between control and patient classes, which can affect supervised and unsupervised classification. We implemented an over-sampling approach known as Synthetic Minority Over-sampling Technique (SMOTE) to correct for this imbalance.^62^ Oversampling was performed using the Themis library in R [65]. This calculates the *k*-nearest neighbors for each observation from the minority class and creates synthetic examples along line segments joining the observations with random nearest neighbors. It has successfully been applied to datasets including electronic health records [63] and image segmentation [64], demonstrating improvements in overall model accuracy. We used SMOTE for supervised classification to augment an imbalanced dataset. SMOTE improves the sensitivity of models tested at the cost of diluting the information of biomarkers in real world sampling. Sub-clusters of depression (subtypes in the minority class) cannot be found using data augmented with SMOTE because the method inherently creates artificial clusters. Synthetic observations are created with values near real observations of the minority class. SMOTE was used with *k* = 3 nearest neighbors to generate data for 19 synthetic patients, resulting in 68 observations total (49 real participants and 19 synthetic observations). This brought the classes to parity. The dataset used for training and tuning included 34 controls and 34 patients. See Supplementary Figure 9 for a visual depiction of the sampling strategy. In Supplementary Figure 9A we depict the augmented dataset created by using SMOTE oversampling. In panel B, we depict the 10 fold cross-validation that was used for model training and hyperparameter tuning. 10 folds of the data are constructed, with each withholding a separate 1/10th of the data for evaluating the model accuracy. This resulted in a stratified sample of the classes (each fold contained an equal number of class observations).

We used the Tidymodels ecosystem in R for training [65]. Model hyperparameter tuning was performed using grid search. Fifty hyperparameter combinations were assessed based on a Latin hypercube assignment. Hyperparameters were chosen based on the model with the highest average Matthew’s Correlation Coefficient (MCC) score across the withheld 10% folds. This method was applied to both the ERP and DCM training data sets separately.

Cross-validation was used to train classifiers and assess whether DCM features can better measure depression’s internal heterogeneity, compared to EEG features.^66^ We compared multiple performance metrics, including *F*-measure (aka F1-score), and the Matthews’ Correlation Coefficient (MCC).^69^ MCC is easier to interpret compared to Cohen’s Kappa, as there is no reference to the expectation of class size, which may not be known and has advantages over the F-measure with regards to class imbalance in binary classification.^70^ Similarly, although hyperparameter search/tuning should usually be conducted on a validation set that is completely separate from training and test sets^71^, here we are comparing the same number of DCM and EEG features, with the same model structure and hyperparameter tuning. Model fitting is only being used to evaluate DCM and ERP’s relative accuracy.

### Feature Importance

The best performing classifier was selected by mean MCC score across all ten folds. This classification algorithm was then used to compute feature importance scores. Shapley additive explanation (SHAP) values were constructed to assess feature importance. SHAP values predictions on the original data set (49 participants, no SMOTE augmentation).^72^ SHAP values were constructed using subsampling of the different combinations of the input features and attributing a weight representing how much credit features should receive for class prediction. SHAP values quantify the importance of a feature based on classification accuracy independently and in conjunction with all possible subsets of features involving the feature of interest (SHAP values arise from scoring the effect of “coalitions” of features). This reveals how efficient the low dimensional space spanned by DCM and EEG features is in describing the internal heterogeneity of patients with depressive symptoms.

In general, SHAP values weight predictive power of features. Feature importance across the entire sample of observations is assessed by adding the prediction weights for all classes. Note that SHAP values can be positive or negative in binary classification (contributing or detracting from a positive classification), therefore absolute SHAP values were used. The predictive power of EEG features vs. DCM features was compared directly because the corresponding SHAP values take on the same scale and are predicting the same underlying data.

### Unsupervised Clustering

The ten most important features as determined by SHAP values from both the ERP and DCM feature sets were used to construct embedding scores with t-stochastic neighbor embeddings (*t-*SNEs). *t-*SNEs are useful for visualizing and exploring higher-dimensional data in lower dimensional representations. *t-*SNEs are scores used for dimensionality reduction when non-linear relationships exist in data.^73^ The method iteratively reduces the KL divergence between a Gaussian probability distribution of similarity between data points in high dimensional space and a *t-*distributed representation in lower dimensional space. Here, we used three dimensions as the lower dimensional space. This provided visualizations of the data that were convenient for assessing subtypes or clusters of patients with depressive symptoms.

Clustering was performed using *k-*means in the three dimensional space obtained by *t-*SNE. *K*-means is a method of unsupervised machine learning that iteratively groups bundles of observations to reduce within-cluster sum squares distances and increase the sum squared distance between cluster centroids.^74, 75^. *K*-means depends on an *a priori* number of clusters. The optimal cluster number can be found by computing Silhouette scores across all candidate values of *k*. Observations which been classified appropriately have a lower mean distance between points within their assigned cluster compared to the mean distance to points in the next-nearest cluster neighbors.^76^ This ratio is given by Silhouette scores. We computed the Silhouette score averaged over all data features (in the low dimensional *t-*SNE space) for all candidate values of *k* from two to twelve.

